# MYC-driven increases in mitochondrial DNA copy number occur early and persist throughout prostatic cancer progression

**DOI:** 10.1101/2023.02.20.529259

**Authors:** Jiayu Chen, Qizhi Zheng, Jessica L. Hicks, Levent Trabzonlu, Busra Ozbek, Tracy Jones, Ajay Vaghasia, Tatianna C. Larman, Rulin Wang, Mark C. Markowski, Sam R. Denmeade, Kenneth J. Pienta, Ralph H. Hruban, Emmanuel S. Antonarakis, Anuj Gupta, Chi V Dang, Srinivasan Yegnasubramanian, Angelo M. De Marzo

**Affiliations:** Department of Pathology, Johns Hopkins University School of Medicine, Baltimore, Maryland, USA; Department of Oncology, Johns Hopkins University School of Medicine, Baltimore, Maryland, USA; Department of Urology, Johns Hopkins University School of Medicine, Baltimore, Maryland, USA; The Sol Goldman Pancreatic Cancer Research Center, Johns Hopkins Medicine, Baltimore, Maryland, USA; Department of Biochemistry and Molecular Biology, Johns Hopkins Bloomberg School of Public Health, Baltimore, Maryland, USA

## Abstract

Increased mitochondrial function may render some cancers vulnerable to mitochondrial inhibitors. Since mitochondrial function is regulated partly by mitochondrial DNA copy number (mtDNAcn), accurate measurements of mtDNAcn could help reveal which cancers are driven by increased mitochondrial function and may be candidates for mitochondrial inhibition. However, prior studies have employed bulk macrodissections that fail to account for cell type-specific or tumor cell heterogeneity in mtDNAcn. These studies have often produced unclear results, particularly in prostate cancer. Herein, we developed a multiplex *in situ* method to spatially quantify cell type specific mtDNAcn. We show that mtDNAcn is increased in luminal cells of high-grade prostatic intraepithelial neoplasia (HGPIN), is increased in prostatic adenocarcinomas (PCa), and is further elevated in metastatic castration-resistant prostate cancer. Increased PCa mtDNAcn was validated by two orthogonal methods and is accompanied by increases in mtRNAs and enzymatic activity. Mechanistically, MYC inhibition in prostate cancer cells decreases mtDNA replication and expression of several mtDNA replication genes, and MYC activation in the mouse prostate leads to increased mtDNA levels in the neoplastic prostate cells. Our *in situ* approach also revealed elevated mtDNAcn in precancerous lesions of the pancreas and colon/rectum, demonstrating generalization across cancer types using clinical tissue samples.

## Introduction

Warburg observed in the 1920s that cancer cells take up excess glucose that is metabolized mostly to lactate, even in the presence of oxygen. He postulated that such aerobic glycolysis (the “Warburg effect”) was universal in cancer, essential for cancer cell proliferation and caused by disrupted mitochondrial respiration (1). The concept of mitochondrial dysfunction as oncogenic was later supported by studies showing that a number of cancer types contain somatic point mutations and/or deletions in mitochondrial DNA (mtDNA) (2–5). In addition, inactivating mutations in nuclear genes whose protein products function in mitochondria occur in several hereditary cancer syndromes, including those caused by mutations in fumarate hydratase (FH) and succinate dehydrogenase (SDH) (6–9). In the prostate, somatic mutations in mtDNA have been proposed as potential cancer drivers (10–18). Furthermore, a 3.4 kb deletion in mtDNA has been reported in prostate cancer, and these and other mtDNA deletions accumulate in the aging prostate (16, 17, 19). Nevertheless, the functional significance of somatic mitochondrial DNA alterations as oncogenic drivers in cancer, including prostate cancer, remains to be fully clarified.

The phenomenon of aerobic glycolysis, which is the use of glycolysis rather than oxidative phosphorylation as a source of energy in cells even in aerobic conditions, was long proposed as a unique property of cancer cells indicative of mitochondrial dysfunction in cancer (20, 21). However, recent evidence has shown that such aerobic glycolysis is not cancer-specific and commonly occurs in non-neoplastic proliferating cells as part of a metabolic program during cell growth and division (1, 20–26). While a complete picture of precisely why dividing cells undergo aerobic glycolysis is still emerging, the metabolism of glucose to lactate helps maintain high levels of glycolytic intermediates and/or NAD^+^ to support anabolic reactions needed for cell growth and replication (1, 21, 22, 24, 27). Furthermore, multiple lines of evidence suggest that cancer and other non-neoplastic proliferating cells have increased mitochondrial activity. Several metabolites generated by the TCA cycle are used as intermediates for macromolecular synthesis in proliferating cells. Also, except for a relatively rare subset of cancer types (e.g., oncocytomas of the kidney, renal tumors associated with inactivation of *FH* or genes encoding the succinate dehydrogenase complex), respiration is not generally markedly impaired in cancer (21, 25, 28), and a functioning electron transport chain is required for proliferation of most normal and cancer cells (21, 29–32). Taken together, it is now apparent that mitochondria display increased activity in most cancers; albeit, their metabolic activities are reprogrammed to facilitate cell proliferation and continuous self-renewal (1, 2, 21, 22, 24–28, 33).

Mitochondrial function is regulated by mitochondrial biogenesis, which is controlled, at least partially, by mtDNA copy number (mtDNAcn) (34, 35). Alterations in mtDNAcn have been reported in cancer (35). For example, increased mtDNA copy number has been described in acute myelogenous leukemia (AML) (36, 37), endometrial carcinoma (38), esophageal squamous cancer (39), pancreatic cancer (5), and colorectal cancer (40). Further, treatment with an inhibitor of mtDNA replication (41) or of a mitochondrial complex 1 inhibitor (42) in AML has shown promise in preclinical studies. Notably, increased mtDNA content was mechanistically linked to microsatellite-stable colorectal cancer cell proliferation and metastasis (40). However, in other cancers, such as gastric, breast, hepatocellular, non-small cell lung and renal carcinomas, decreased mtDNA levels, as compared to their normal tissue counterparts, have been reported (35). Thus, there is no general pattern regarding mtDNAcn changes in tumor-normal comparisons in common adult cancer types.

In terms of mtDNAcn changes in prostate cancer, there are contradictory results (16, 43–45). Mizumachi et al. examined single laser microdissected cells from a relatively small number (N=9) of prostate cancers and reported increased mtDNA content in some cancers, decreased amounts in others, and a general increase in cell-to-cell variability in cancer cells compared to matched benign tissues (43). Koochekpour et al. reported lower levels of mtDNA in human prostate cancer compared to normal prostate tissues and suggested that reduced levels of mtDNA can drive aggressive prostate cancer (44). Reznik et al. analyzed mtDNAcn variation across multiple cancers relative to matched non-neoplastic tissues using next generation sequencing (NGS) data from The Cancer Genome Atlas (TCGA). This study found that prostate cancer had non-statistically significantly increased bulk tumor mtDNA levels compared to benign prostate tissues (45). Schöpf et al. reported that there was no significant difference in mtDNAcn between benign and malignant prostate tissues, although mtDNAcn was increased in higher grade cancers (18). Higuchi et al. showed that androgen-independent C4-2B cells had lower levels of mtDNA than androgen-sensitive LNCaP cells and that depletion of mtDNA in LNCaP cells resulted in the development of androgen independence (46). Consequently, whether there are increased or decreased levels of mtDNA in prostate cancer, and whether mtDNA levels are associated with disease aggressiveness and castration resistance remains unclear. To improve our understanding of the role of mtDNAcn alterations in prostate cancer, it is important to resolve these inconsistent findings.

A concern with most prior studies of mtDNAcn alterations in cancer is that they usually employ bulk macrodissection of tumor and matched benign tissues, which is followed by cellular disruption and DNA isolation. These bulk mtDNA measurements could not take into account heterogeneity within or between tumor nodules or the diversity of cell types (47). We have shown, using a novel *in situ* hybridization approach, differing mtDNA levels in normal tissue compartments and cell types in most organs (48). Therefore, the use of bulk macrodissections may be ill-suited for tumor/normal comparisons by not accurately representing differences among relevant neoplastic and non-neoplastic cell types or regional heterogeneity in mtDNA levels of neoplastic cells (43). To understand better the potential biological role of mtDNAcn changes in cancer initiation and progression, it is critical to determine when during the carcinogenesis process the alterations occur (49). Since cancer precursor lesions are often difficult to obtain from fresh tissues, and are often microscopic in size, most studies of mtDNAcn alterations on such precursors require either laser capture microdissection (LCM) or an *in situ* approach (49).

Here, we report the enhancement of our *in situ* approach to mtDNA quantification (48) by combining it with iterative sequential immunohistochemistry in a multiplex assay, allowing for cell type and compartment specific mtDNA measurements. This multiplex assay revealed generally increased mtDNAcn in pre-neoplastic lesions and prostate cancer. Because we and others have linked MYC overexpression to prostate cancer development and progression (50–52), and because MYC has been shown to induce mitochondrial biogenesis (53), we also investigated whether MYC is a key factor regulating mtDNA copy number in human and murine prostate cancer cells. These studies have revealed that MYC increases mtDNAcn through stimulating mtDNA replication apparently by increasing expression of mtDNA replication factors, including the helicase TWINKLE.

## Methods

### Mice

Hi-MYC mice (54) on an FVB/N background (n = 3, age = 2-6 months) were conventionally housed according to protocols approved by the Animal Care and Use Committee at the Johns Hopkins University School of Medicine. Animal tissues were harvested and fixed in 10% Neutral buffered formalin for 48 hours.

### Chromogenic mtDNA in situ hybridization combined with immunohistochemistry

*In situ* hybridization for mtDNA for both fresh frozen tissues and formalin-fixed paraffin-embedded (FFPE) tissues was performed based on a previous method (48). The tissue pretreatment conditions for human prostates was as previously described (48). For human colon FFPE tissues, the pretreatment was the same as that for human prostate FFPE tissues (48). For human pancreas FFPE tissues, the pretreatment was 15 min steaming in RNAscope® Target Retrieval reagent followed by 10 min incubation in protease plus at 40 °C.

The mtDNA *in situ* hybridization signals were developed using 3-amino-9-ethylcarbazole (AEC, Vector Laboratories, SK-4200), counterstained in hematoxylin and blued in Shandon bluing buffer (Thermo Scientific, 6769001). The slides were mounted, coverslipped, scanned, and de-coverslipped. The AEC was de-stained in an alcohol gradient. The slides were then rehydrated in an alcohol gradient and incubated using a HRP blocker (ACD bio) for 15 min at 40 °C before being subjected to sequential immunohistochemistry. This scanning, destaining and blocking process was repeated between all staining cycles.

After *in situ* hybridization, the slides were incubated with an anti-CK8 antibody (1:1500, Abcam, ab53280) for 45 min, PowerVision Poly-HRP anti-Rabbit IgG (Leica, PV6114) for 30 min, and visualized by incubation in AEC for 30 min, followed by counterstaining and bluing. For some slides, an additional CK903 IHC staining cycle was performed after CK8 detection. CK903 (1:50, Enzo, ENZ-C34903) was used as the primary antibody, PowerVision Poly-HRP anti-Mouse IgG (Leica, PV6119) as the secondary antibody, and AEC as the chromagen. All primary antibodies were diluted in Antibody Dilution Buffer (Ventana, adb250) unless specified.

The mtDNA *in situ* hybridization on mouse prostate tissues using *Mm-mt-Co1* sense probe was performed as described previously (48). *Hs-MYC* mRNA *in situ* hybridization was performed similarly with probes against human *MYC* (ACD Bio, 311761).

### Multiplex Fluorescent In Situ Hybridization and Immunofluorescence

Multiplex fluorescent *in situ* hybridization for mitochondrial RNA(s) expression was performed using HiPlex12 detection kit (ACD Bio, 324105) on fresh frozen tissues. Briefly, fresh frozen prostate tissues were hybridized with RNAscope HiPlex probes against *Hs-MT-CO1* sense (mtDNAcn), *Hs-MT-CYB*, *Hs-MT-ATP6* and *Hs-MT-RNR1* in two rounds. Whole slides were scanned and fluorophores were cleaved between rounds. After HiPlex, the slides were then incubated in anti-CK8 and anti-CK903 antibodies with the same concentration as above. Secondary antibodies were the same except they were used at 1:125 dilution. Lastly, the slides were developed with Opal fluorescent dye at 1:50 dilution (Akoya Biosciences) in different channels. Catalog numbers for HiPlex probes were the following: *Hs-MT-CO1* sense (478051), *Hs-MT-CYB* (582771), *Hs-MT-ATP6* (532961) and *Hs-MT-RNR1* (425961), all from ACD Bio.

### Immunohistochemistry (IHC)

All single-plex 3,3′-Diaminobenzidine (DAB) based chromogenic IHC on tissue samples (FFPE or fresh frozen) was performed on an automated Ventana Discovery ULTRA system (Roche). Fresh frozen tissues were air dried at room temperature for 15 minutes, fixed overnight in 10% neutral buffered formalin and then rinsed off with dH2O prior to staining. IHC labeling for c-Myc was performed using the DISCOVERY HQ+ Amp kit with primary antibody against c-MYC (1:600, Abcam, ab32072). IHC for EdU and MYC on cells grown on chamber slides was performed by following the protocol described before (48). A primary antibody against BrdU (Abcam, ab136650) was first validated to be specific for EdU as well, and used at 1: 1000 dilution. The primary antibody against c-MYC was used at 1:250 dilution (Abcam, ab32072). The secondary antibodies were PowerVision Poly-HRP anti-rabbit or anti-mouse (Leica, PV6114 and PV6119) and developed with DAB.

### Quantitative PCR

Quantitative PCR to measure mtDNAcn from DNA extracted from LCM prostate samples was conducted as described previously (48). Probes used in this study were *Hs-MT-CO1* (Thermo Fisher, Hs4331182) as the mitochondrial gene and *BGLT3* (Thermo Fisher, Hs01629437) as the nuclear gene reference.

### Image Analysis

Whole slides were scanned at x40 resolution (0.25 um per pixel) on a Ventana DP200 slide scanner (Roche) for chromogenic slides, or a TissueFAXS slide scanner for fluorescent slides (TissueGnostics). All the image processing and analysis were performed using HALO version 3.0 (Indica labs). Single-staining whole slide images in the chromogenic multiplex *in situ* hybridization and immunohistochemistry assays were fused using the Serial Staining Image Fusion protocol provided by HALO as described (55). Briefly, after scanning, each set of images were registered. Images were color deconvoluted in a case-by-case manner based on red, green, blue (RGB) optical density values for each marker in AEC and for hematoxylin. During this process, the registration transforms generated in the previous step were used to align all slides into the same coordinate system. The pseudocolor single stained whole slide images were then fused with the same image resolution as the original images, and then subjected to downstream analyses. Fluorescent scanning images were exported using TissueFAXS viewer software (TissueGnostics) in single channel images and fused similarly in HALO.

A random forest algorithm in the Tissue Classifier Add-on of HALO was used to distinguish epithelium from stroma and empty tissue compartments. The calculated area of each compartment type was based on the CK8 and hematoxylin status for each slide. CK8 was chosen as it is positive for all normal prostatic cells, prostatic intraepithelial neoplasia (PIN), primary and metastatic prostate cancer. In needle biopsies from metastatic sites in the liver, in which hepatocytes are CK8-positive, the metastatic carcinoma regions were manually outlined. For samples that were also stained with CK903, the classifier was additionally trained to recognize and calculate the area of prostate basal epithelium vs. luminal epithelium. Since there were slide-to-slide differences in CK8 and CK903 intensity, classifiers were trained for each slide individually. All classifiers had resolution set at 1.04 μm/px and minimum object size at 20 μm^2^. For radical prostatectomy samples, regions of normal prostatic epithelium, HGPIN and prostate cancer were annotated before analysis. To quantify mtDNA *in situ* hybridization signals, the Area Quantification FL algorithm (v.2.1.2) and above-mentioned classifiers were used in combination, and percentages of mtDNA signal area (%mtDNA area) in each classified compartment (e.g. prostate basal cells, prostate luminal cells, or total prostate epithelium [basal + luminal cells]) were measured and reported. We have previously shown that this %mtDNA area is strongly correlated with mtDNAcn, as defined by the ratio of mtDNA levels to nuclear DNA levels (48). The quantification algorithm of each image was individually optimized using the real-time tuning function based on both the deconvoluted image and the original chromogenic image.

### Cell lines and cell culture

PC3, LNCaP, CWR22Rv1 prostate cancer cells were cultured in RPMI1640 medium supplemented with 10% (v/v) fetal bovine serum (FBS) at 37°C and 5% CO2. Cells were maintained on cell culture flasks (Sarstedt) or CC2 chamber slides (Thermo Scientific, 154739). MYCi975 was originally reported by Han et al. (56) and was obtained from AxonMedChem (Axon 3229).

### Western blotting

Whole cell extracts from cell lines were obtained using RIPA buffer (Thermofisher) containing a protease inhibitor cocktail (Roche) and benzonase (Thermofisher). Protein concentrations were quantified using a BCA gold assay (Thermofisher). Equal amounts of total protein were prepared in 4X NuPAGE LDS Sample Buffer (NP0007, Invitrogen) and 5% BME, and boiled at 95°C for 5 min. The protein samples were resolved on 4-12% NuPAGE Bis-Tris gel with protein molecular weight standards (PageRuler Prestained Protein Ladder, 10 to 180 kDa, Thermofisher). The gels were transferred onto PVDF membranes using the Trans-Blot Turbo Transfer System (Bio-Rad). Primary antibodies used in this study included: rabbit anti-c-Myc antibody [Y69] (ab32072, Abcam, 1:1000), mouse anti-β-Actin antibody (3700, Cell Signaling, 1:5000). Fluorescent secondary antibodies used in this study were IRDye 680RD Donkey anti-Mouse IgG Secondary Antibody and IRDye 800CW Donkey anti-Rabbit IgG Secondary Antibody (Licor, 1:5000).

### COX and SDH enzyme histochemistry

COX and SDH enzyme histochemistry was performed on fresh frozen human prostate tissues. Briefly, fresh prostate tissues were collected, frozen, cryosectioned into 5 μm (SDH) or 10 μm (COX) sections. The frozen slides were air dried at room temperature for 1 hour. COX reaction mix containing 25 µL of 4 mM Cytochrome c, 67 µL of 10 mg/mL 3,3′-diaminobenzidine tetrahydrochloride (DAB) in 1 mL 1X PBS was freshly prepared, followed by addition of 2 µg bovine catalase and vortexing, and added to slides for 40 min at 37 °C in a moist chamber. SDH reaction mix was prepared using 1.5 mM NBT, 130 mM sodium succinate, 0.2 mM PMS, and 1.0 mM sodium azide in 1X PBS. The mixture was applied to tissue slides for 40 min at 37 °C in a moist chamber. After either COX or SDH reaction was finished, the slides were washed in PBS for 4 x 10 min, dehydrated in an ethanol gradient and xylene, mounted and coverslipped in Cytoseal 60.

### EdU-immunoprecipitated DNA and real time quantitative PCR

To measure newly synthesized mtDNA, we developed a method for EdU-immunoprecipitated DNA and real-time qPCR based on a similar approach used by Jiang et al. (57). Briefly, cells grown on flasks were treated with MYCi975 at the indicated concentrations for 48-96 hours. Before harvesting, the cells were incubated in cell culture media containing 10 µM EdU for 4 hours. Cell pellets were divided into two portions, one for total DNA extraction and the other for total protein extraction. Total DNA was extracted using DNeasy blood and tissue kit (Qiagen). Immunoprecipitation of EdU-containing DNA was performed using the iDeal ChIP-seq kit for Transcription Factors (Diagenode) with the following modifications. For each immunoprecipitation (IP) reaction, 30 μL of washed Protein A-coated magnetic beads was combined with 70 μL IP reaction mix containing 6 µL of BSA, 20 µL 5x iC1b buffer, 0.5 µg antibody in ChIP grade water and incubated for 2 hours at 4°C under rotation. Equal amounts of total DNA (5-15 µg) were added to the reaction mix and incubated overnight at 4°C under rotation. The beads were washed and the DNA was eluted and purified by following the manufacturer’s protocol. One percent of the INPUT was subjected to the same elution and purification process. DNA (1 μL) from the purified IP and INPUT samples was used for real time quantitative PCR. TaqMan Universal PCR Master Mix II (no UNG) and TaqMan probes (mitochondrial: *MT-RNR1*, *MT-CO1* and nuclear: *BGLT3*) were used following the manufacturer’s protocol (Thermofisher). EdU-containing newly synthesized DNA was calculated and reported by comparing the relative amount of immunoprecipitated DNA to the INPUT DNA for specific mitochondrial and nuclear genes (% of recovery). The anti-BrdU/EdU antibody (Abcam, ab136650) was used, and Mouse (G3A1) mAb IgG1 (Cell Signaling, 5415) was used as the negative isotype-matched control for the IP reactions.

### mtDNA copy number and RNAseq in laser capture microdissected prostate tissues

Fresh frozen tissues were placed in OCT and sections were cut at 7 μm thickness and mounted onto membrane slides (Leica, 11600288). The slides were fixed in ice-cold 70% EtOH for 5 min, hydrated in RNAse free water, counterstained in hematoxylin for 20 seconds, washed twice in RNAse-free water, and air dried. The slides were stored at 4°C for a short time before microdissection. LCM of regions of interest was performed using Leica LMD 7000 Microscope. Tissue digestion and DNA/RNA extraction was performed using Allprep DNA/RNA Kits following the manufacturer’s recommendations (Qiagen, 802804). For DNA library preparation, samples were sheared by sonication on the Covaris S2 System; libraries were constructed according to the protocol provided in the Illumina Nano TruSeq Library prep guide. Assessment of the yield and size distribution of the amplified library was performed on the Agilent 2100 Bioanalyzer using the High Sensitivity Chip. Libraries were then pooled at equal molar concentrations for sequencing. Sequencing was performed using Illumina HiSeqX and NovaSeq6000 with Paired End 150 bp x 150 bp read configuration. For sequencing data processing, trimgalore v0.6.3 was used to trim the reads. Bwa v0.7.7 (mem) was used to align to the hg19 and hg38 human genome builds. Piccard-tools v1.119 & GATK v3.6.0 were used to create a recalibrated bam file. Bedtools v2.27.1 (genomecov) was used to determine the nuclear coverage (i.e., all chromosomes except mtDNA) and the mitochondrial coverage. mtDNA copy number was computed using the following formula: mitochondrial coverage/nuclear coverage. RNAseq and gene expression measures on mtDNA replication related genes were performed as previously described (58). The analysis of the WGS and RNA-seq data here is limited to just the reported mtDNA copy number assessment and the gene expression measures for known mtDNA replication genes. Full details and reporting of the WGS and RNA-seq data will be reported as part of another study.

### Gene set enrichment analysis

Gene set enrichment analysis on microarray data from cell lines with MYC knockdown (59) was performed using the fgsea R package (60). The mtDNA replication related gene set was obtained from the Mitocarta 3.0 database (61).

### Public data analysis

Publicly available ChIP-seq and RNA-seq data on cells treated with MYCi975 were obtained from NCBI GEO accession number GSM5399497 GSM5399571, GSM5399627, and GSM5399631 (62) and visualized using the Integrated Genomics Viewer (IGV). RNA-seq data on *MYC* and *TWNK* in different cancer types from TCGA PanCancer Atlas was accessed using cBioportal (63, 64).

### Statistics

Statistical analysis and visualization was performed using Prism GraphPad 9.0 and R version 4.0.2. Image analysis results between different lesion types were compared using nonparametric Mann Whitney test.

### Study approval

*Human studies.* Human prostate tissues (frozen, or formalin fixed and paraffin embedded [FFPE]) were obtained from radical prostatectomies performed at The Johns Hopkins Hospital. Their use was approved by the Johns Hopkins University School of Medicine Institutional Review Board (NA_00087094). **Supplementary Table 1** summarizes the clinical features of the cases used for *in situ* hybridization study. **Supplementary Table 2** summarizes the clinical features of the patients participating in the whole genome sequencing study. *Animal studies.* All animal study protocols were approved by the Animal Care and Use Committee at Johns Hopkins University School of Medicine.

## Results

### Basal cells show higher mtDNAcn than luminal cells in benign prostate glands

Using a mtDNA chromogenic *in situ* hybridization (CISH) assay, we previously observed that mtDNA signals were visually higher in prostate basal cells compared with normal luminal cells (48). To quantitatively assess mtDNAcn on tissues, we combined mtDNA CISH with multiplex immunohistochemistry (IHC) using two protein markers, keratin 8 (CK8) and a combination of keratins 5+14 (CK903). Anti-CK8 labels both basal and luminal prostate epithelial cells, and, in normal (e.g. non-atrophic) epithelium, anti-keratins 5+14 labels only prostate basal cells. We employed an iterative chromogenic approach (55) in which we first performed *in situ* hybridization for mtDNA followed by whole slide scanning which was then followed by sequential IHC for each of the keratin antibodies (**Fig. 1**). We used radical prostatectomy specimens to compare regions containing normal appearing epithelium, HGPIN and adenocarcinoma. **Fig. 1** shows the pseudocolor image and the cellular compartmental segmentations from a representative case containing adjacent normal appearing epithelium and invasive adenocarcinoma. Quantitative image analysis demonstrated that basal cells contained an approximately 5-fold higher mtDNA percent signal area on average than normal appearing luminal cells (**Supplemental Fig. 1**).

**Figure 1.**
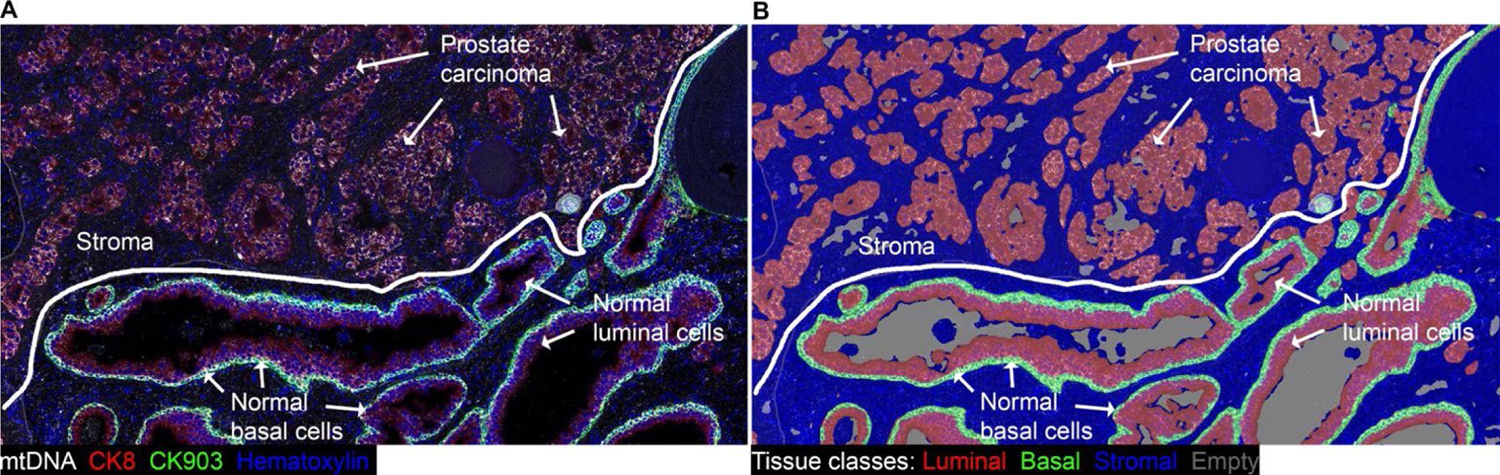
Multiplex chromogenic *in situ* hybridization for mtDNA and IHC in human prostate. (**A**) Whole slide scans of each staining round were imported into HALO and registered followed by color deconvolution, scan fusions and application of pseudocolors to the resultant channels. mtDNA *in situ* hybridization signals are shown in white, epithelial cells (both basal and luminal) stained with CK8 in red, basal cells stained with CK903 in green and nuclei in blue (hematoxylin channel). (**B**) Markup image of classifier result showing basal cells (green, present in normal), luminal cells (red, present in normal and at all stages of prostate cancer tumorigenesis), stroma (blue) and empty space/lumens (gray). Original magnification, x120.

### Increased mtDNAcn in high grade prostatic intraepithelial neoplasia (HGPIN)

HGPIN is the likely precursor to most human prostate cancers (65, 66), yet mtDNAcn levels have not been studied in these lesions. We sought to compare neoplastic epithelial cells in HGPIN to matched benign normal appearing epithelium. By visual inspection, luminal cells in HGPIN showed increased mtDNA signals as compared with normal luminal cells (**Fig. 2A**). By image analysis, HGPIN showed a consistent increase in the median percent mtDNA signal area of approximately 3-fold compared to the normal prostate glands (**Fig. 2F**). This increase was found despite the fact that basal cells were not excluded from the image analysis in the normal regions.

**Figure 2.**
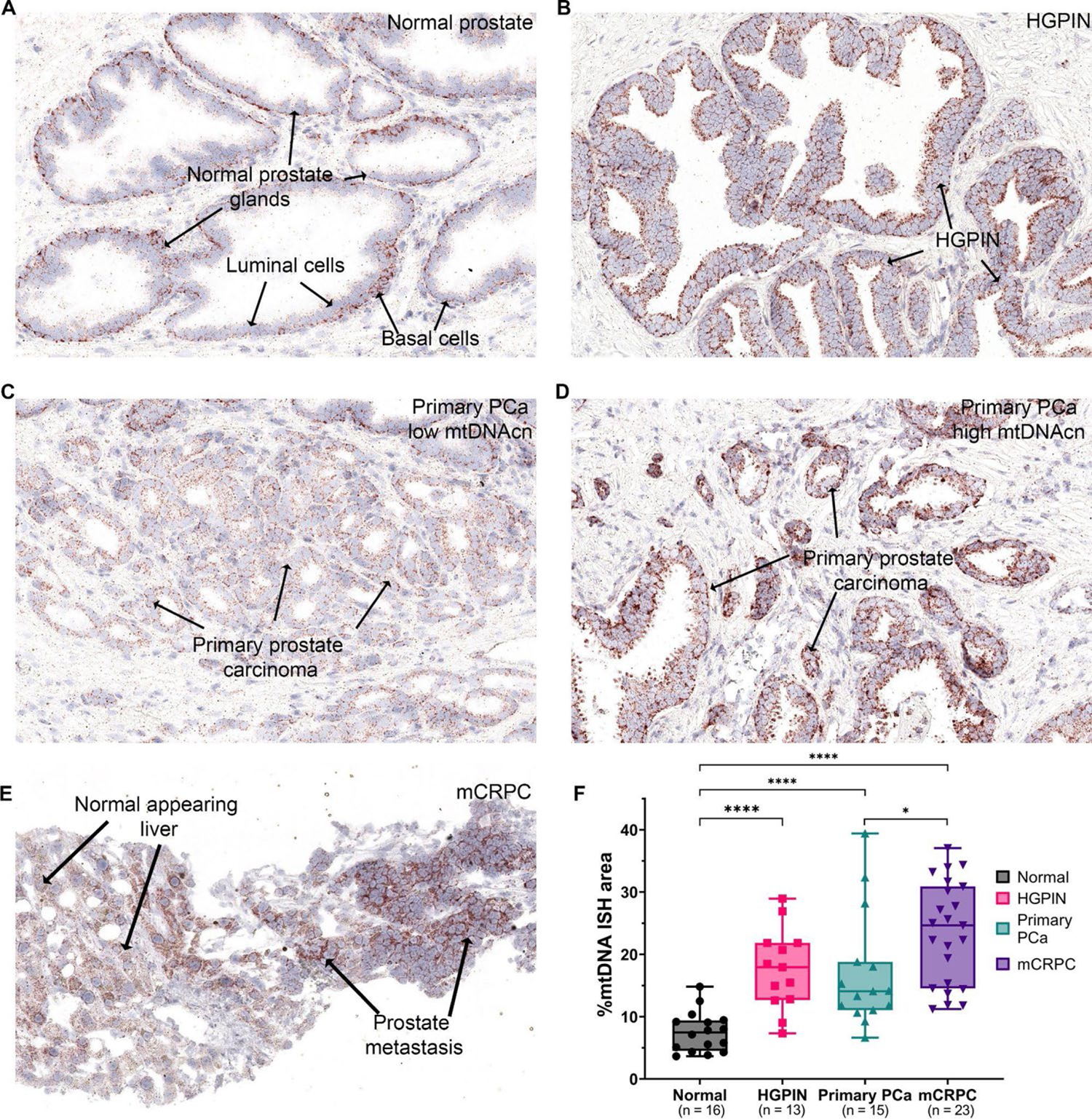
Chromogenic *in situ* ybridization for mtDNA from prostate samples at different stages of prostatic tumorigenesis and image analysis results. Tissues were subjected to multiplex CISH-IHC as in Fig. 1, and images from the round of mtDNA *in situ* hybridization are shown. (**A**) Normal appearing prostate glands (**B**) HGPIN, (**C**) invasive adenocarcinoma with low mtDNA CISH signals, (**D**) invasive adenocarcinoma with high mtDNA CISH signals, (**E**) metastatic prostate adenocarcinoma to the liver. (**F**) Quantification of mtDNA in epithelium presented as %mtDNA area at each stage. Each symbol in the graph is the average of %mtDNA area from multiple regions in a single patient. Note that images shown in A-D were from the same patient. Original magnification, x200 (**A-E**). *P < 0.0332, and ****P < 0.0001.

### Increased overall and heterogeneously distributed mtDNAcn in primary prostate cancer

Using the same tissue specimens, we next examined primary invasive adenocarcinomas. Visually, invasive carcinomas generally showed higher mtDNAcn than matched normal regions. Interestingly, there was considerable regional heterogeneity of mtDNA *in situ* hybridization signals, both within individual tumor foci (intratumorally) and between foci separated in space (intertumorally) (**Fig. 2C-D**; **Supplemental Fig. 3**). Quantitative analysis including the different subregions of tumor foci is shown in **Supplemental Fig. 2**. As an example of intratumoral heterogeneity in a primary prostatic adenocarcinoma, **Sup. Fig. 3** shows striking mtDNAcn heterogeneity within an individual tumor from a prostatectomy case. In addition to being contiguous in space, this tumor showed homogeneous ERG expression, indicative of a *TMPRSS2-ERG* gene rearrangement (67, 68). While additional analyses would be needed to definitively assess clonality, this homogeneous ERG expression is consistent with the likelihood that this large tumor with heterogeneous mtDNAcn represents a single clonal origin (**Supplemental Fig. 3**) Taken together, carcinoma lesions had significantly higher levels than normal epithelium, and were similar to HGPIN (**Fig. 2F**).

**Figure 3.**
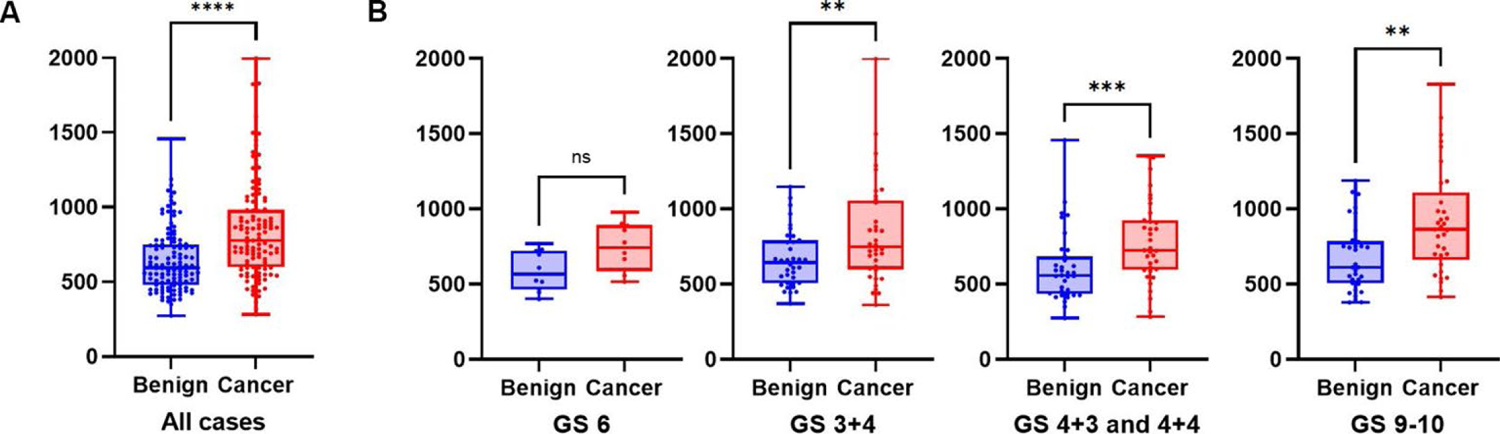
Increased mtDNAcn in prostate cancer using whole genome sequencing. (**A**) Combined dot and box and whisker plot showing increased mtDNA in tumor samples; P<0.0001 by paired t-test for tumor vs. normal. (**B**) Increased mtDNAcn was present across all Gleason grade groups (GG; GS 3+3=GG1, GS 3+4=GG2, GS 4+3 = GG3 and 4+4 = GG4; GS9-10=GG5). Note that GG3 and GG4 are combined due to the relatively low number of GG4 (4+4=8) cases. **P < 0.0021, ***P < 0.0002, and ****P < 0.0001. Blue dots represent benign samples, and red dots represent cancer samples.

As an independent method for comparing mtDNAcn in prostatic tumor-normal pairs, we used whole genome sequencing from laser capture microdissected tissues in which tumor regions and matched benign paired prostatic regions (non-high grade PIN containing) were isolated separately (N=115 prostate cancer cases). In this cohort, which is part of a comprehensive genomic analysis of prostatectomy samples, tumor samples showed increased mtDNAcn compared with matched normal appearing glands (**Fig. 3A**). **Figure 3B** shows the increased mtDNA levels in carcinomas occurred across all major Gleason Grade groups. The distribution of cases by grade, pathological stage, race and other demographic features are shown in **Supplementary Table 2**. While there was no clear increase in mtDNAcn by grade, all of the cancers with mtDNAcn signals above 1000 were Gleason grade group 2 or higher.

A prior study has reported that tumor and benign prostate tissues contain differing levels of mtDNAcn in Black compared to White patients (44). When comparing mtDNAcn in tumor samples between White and Black men (self-identified race) using our NGS approach, there were highly similar mtDNAcn levels between the two, with no statistical difference (**Supplemental Fig. 4A**).

**Figure 4.**
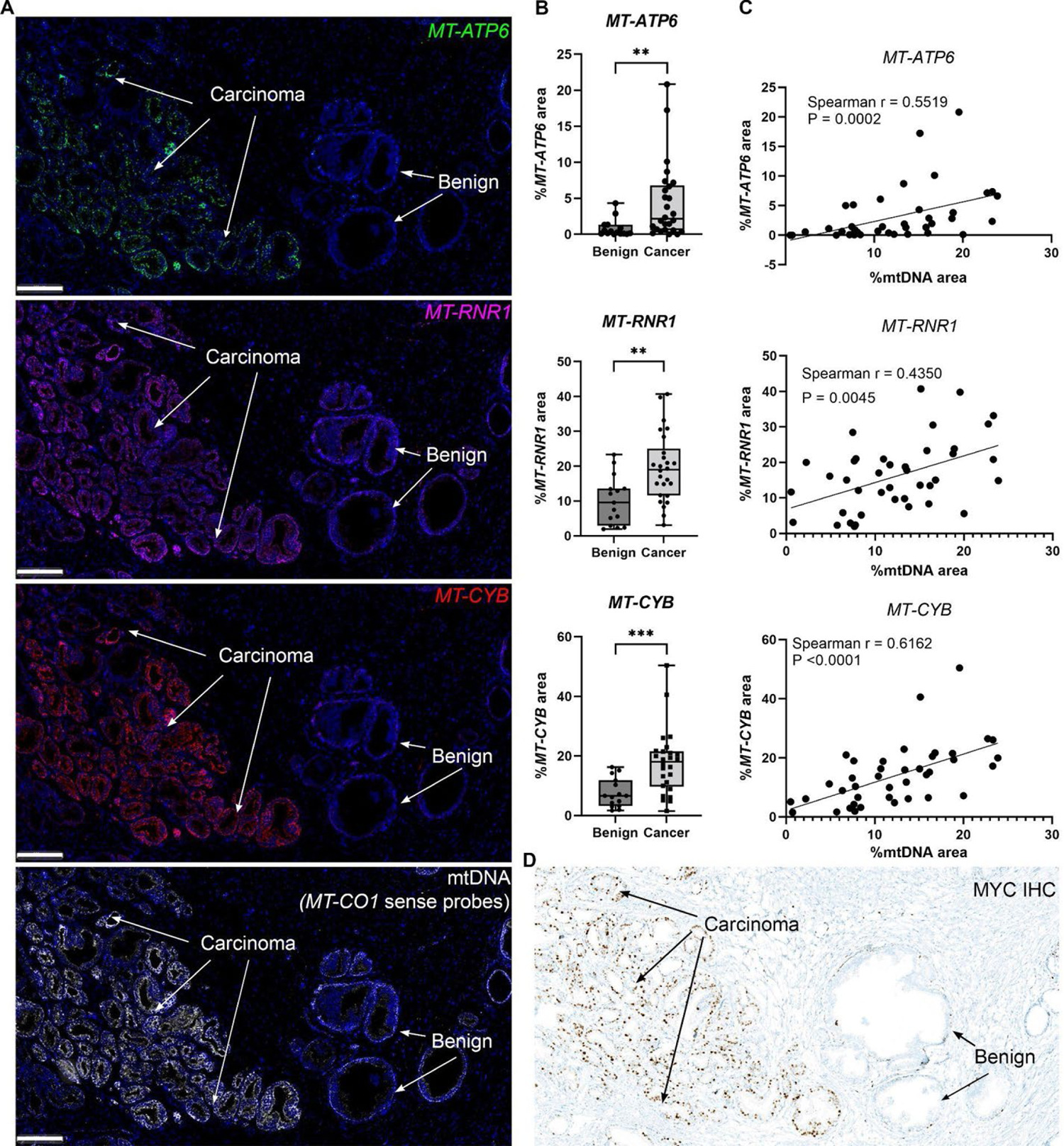
Increased mtDNAcn was accompanied by elevated mtRNA expression as well as MYC protein expression in the prostate. (**A**) Representative images of simultaneous multiplex detection and visualization of mtDNA and mtRNA(s). (**B**) Image analysis results of 3 mtRNA(s) expression in cancer regions compared to the adjacent benign tissues. Each symbol represents an analyzed region of interest. (**C**) Correlations between mtDNAcn and mtRNA expression by image analysis. (**D**) IHC for MYC from an adjacent frozen section of prostate tissue from the same tissue shown in **A**. Original magnification, ∼x70. **P < 0.0021 and ***P < 0.0002. Scale bar = 200 μm (**A**).

As an orthogonal validation of the mtDNAcn estimates using our NGS samples and the analysis pipeline, aliquots of the same DNA samples used for WGS were subjected to quantitative PCR, as described previously (48). This showed a strong positive correlation between the mtDNAcn obtained using WGS and qPCR (**Supplemental Fig. 4B**), validating the quantitative nature of our NGS approach. Also, qPCR on a subset of the samples used for WGS also showed an increase in mtDNAcn in matched tumor samples compared with benign (**Supplemental Fig. 4C**). Thus, three independent methods used in this study all show increased mtDNAcn in tumors versus normal prostatic tissues.

### Metastatic castration-resistant prostate cancer (CRPC) showed higher mtDNAcn with less heterogeneity than the primary prostate cancer

Next, to assess mtDNAcn in advanced prostate cancer, we performed the same iterative CISH-IHC and image analysis workflow using biopsies from men with known metastatic castration-resistant prostate cancer (mCRPC) (N=23 patients). The mCRPC samples were from pretreatment biopsies of patients in two different clinical trials (51, 69). Visually, these metastases demonstrated high mtDNA *in situ* hybridization signals compared with the adjacent non-neoplastic tissues (when present), with some cases showing higher mtDNA signals than hepatocytes in liver, which are known for high mtDNA content (**Fig. 2E**) (48). By image analysis, we observed high mtDNAcn in most of these mCRPC biopsies, which appeared higher as a group when compared to the primary tumors (**Fig. 2F**).

### Functional relevance of increased mtDNAcn in prostate cancer

To address the functional significance of mtDNA alterations in prostate lesions, we first sought to determine whether the increased mtDNAcn was accompanied by corresponding increases in mtRNA(s). We used a multiplex fluorescent *in situ* hybridization assay to perform simultaneous hybridizations for mtDNA along with mtRNAs from three different mtDNA-encoded genes. For the mtRNAs, in the normal appearing prostate regions, we found generally increased signals in the basal versus luminal compartments, and in carcinoma tissues there were increased levels for all 3 of these mtRNAs (**Fig. 4A-B**). Furthermore, there was a correlation between each of these 3 mtRNA species and mtDNA (*MT-CO1* sense) (**Fig. 4C**). In fact, the mtDNA levels correlate with the levels of both MT-rRNA and MT-mRNAs that it encodes. These observations suggest that mtRNA levels may be regulated, at least in part, by mtDNA levels. In addition, we performed enzyme histochemistry for cytochrome c oxidase (COX) and succinate dehydrogenase (SDH) on frozen sections from a number of cases. In general, there were visually higher levels of both enzyme activities in basal cells compared to luminal cells (**Sup Fig. 5**). In prostate carcinomas, there were visually apparent increases as well as heterogeneity that were similar in pattern to the mtDNA *in situ* signals from adjacent tissue sections, providing further support for the hypothesis that the increased mtDNAcn is functional with respect to production of the encoded genes. Interestingly, the lesions with high mtDNAcn, mtRNA expression and mitochondrial enzymatic activities also showed high MYC protein expression, suggesting a mechanistic link between them (**Fig. 4D**).

**Figure 5.**
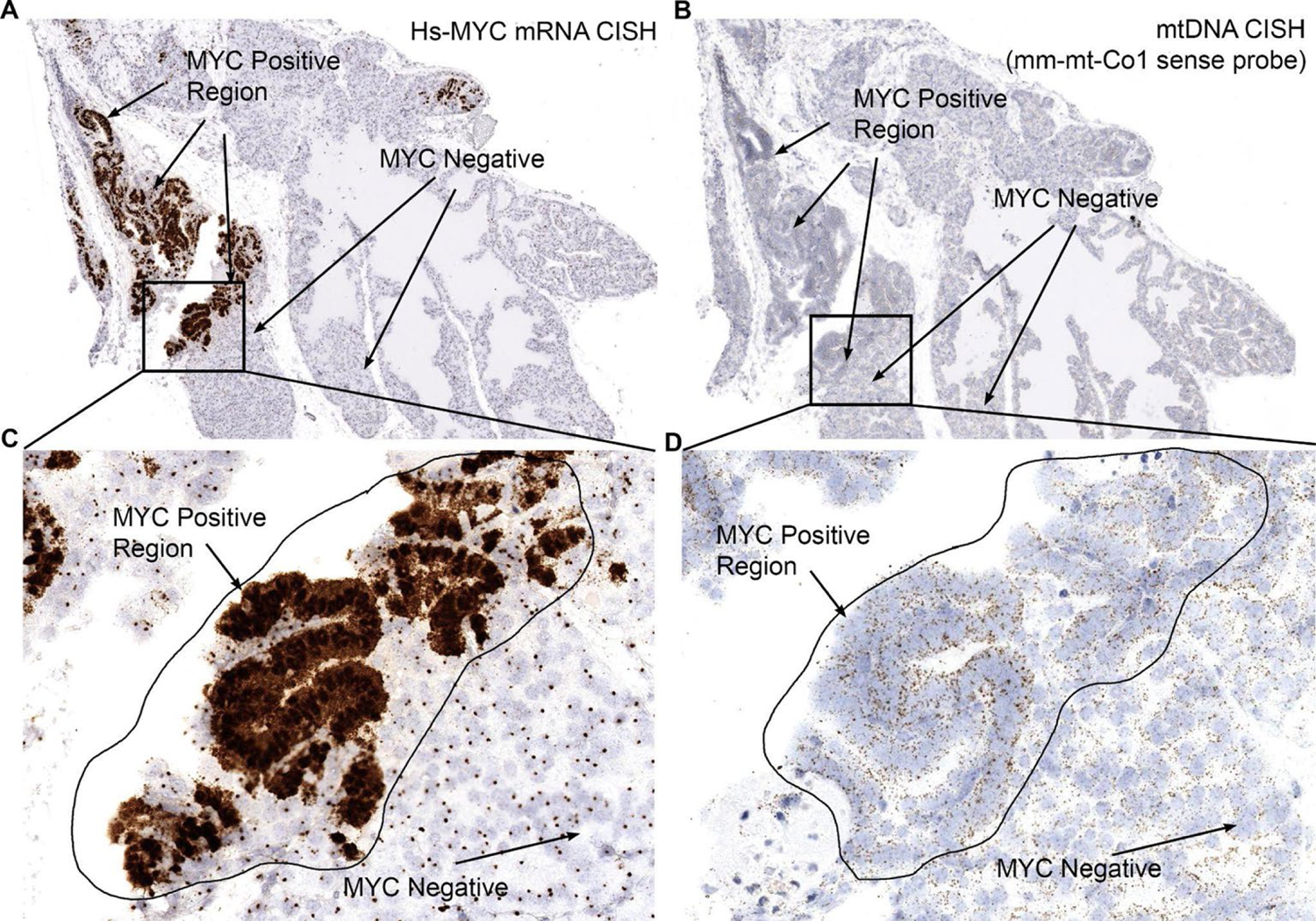
Mouse mtDNAcn is increased in prostates of the Hi-MYC mouse model. Through chromogenic *in situ* hybridization, human MYC mRNA (**A** and **C**) and mtDNA levels (**B** and **D**) were visualized (positive signals are brown). Note the tight spatial correlation with the appearance of cytoplasmic *MYC* signals, corresponding to human *MYC* mRNA expression, and increased mtDNA signals. Also note that the *MYC* probe hybridizes to all nuclei in the transgenic mice as a result of binding to the transgene present in all of the Hi-MYC mouse cells. Original magnification, ∼x70 (**A** and **B**), x200 (**C** and **D**).

### MYC as a Driver of mtDNAcn in Clinical Prostate Cancer

To begin to assess the molecular mechanism for increased mtDNA levels in HGPIN and PCa, we focused on MYC, since MYC has been shown to regulate mitochondrial biogenesis and mtDNA levels (53, 70). MYC is also a key driver of prostate cancer, whose overexpression starts very early in the disease process in HGPIN (66, 71, 72), continues in primary carcinomas and is also highly expressed in mCRPC (52, 71–73). In human prostate cancer cell lines, we have shown that forced overexpression of MYC results in increased mtDNA (48). Despite these findings, since some MYC-driven processes are context and cell type-specific, including its effects on mitochondrial related gene expression (53, 70), it is important to investigate whether MYC could be a driver of mtDNAcn changes in normal prostatic epithelial cells *in vivo* in the context of intact tissues. Therefore, we examined mice with targeted MYC overexpression in the prostate in which MYC is known to drive the development of PIN lesions (Hi-MYC mice) (54, 74, 75). We focused on the anterior prostate lobe of Hi-MYC, since, despite being driven by AR signaling, MYC expression is quite heterogeneous and it often contains regions of MYC-positive and MYC-negative epithelium within the same lobe. We observed that in regions showing increased MYC mRNA and protein levels, there were increased levels of mouse mtDNA that correspond to the regions of MYC overexpression and accompanying morphological changes diagnostic of PIN, including marked nucleolar enlargement (4 of 4 regions in 3 mice). **Fig. 5** shows a representative case in a Hi-MYC mouse.

### Decreased mtDNAcn in response to high dose testosterone treatment in mCRPC and correlation with MYC

Prior work has shown that the cyclical treatment of men with mCRPC with supraphysiological levels of testosterone (SPA, or supraphysiological androgen), referred to as Bipolar Androgen Therapy (BAT), produces tumor regression and clinical benefit in ∼30-40% of patients (51, 76–80). Many of the samples from men with mCRPC shown in **Fig. 2** were from the pretreatment time point of a recently conducted trial, referred to as the COMBAT-CRPC study (51) (clinicaltrials.gov NCT03554317). In this trial, patients received a pre-treatment biopsy of an accessible metastatic site, as well as a biopsy after 3 cycles of SPA (timepoint referred to as cycle 4, day 1 or C4D1). In COMBAT-CRPC, there was a similar response rate (∼40%) to previous reports. Interestingly, most patients in the trial who responded to SPA also showed a marked reduction in MYC protein expression, by quantitative IHC, as well as MYC mRNA by RNAseq. This was accompanied by reductions in Ki67 protein as well as a number of proliferation associated mRNAs (51, 81). We performed the mtDNAcn CISH-IHC assay on all available mCRPC patient samples that had matched post-treatment biopsies from COMBAT-CRPC. **Fig. 6** shows that there was a decrease in mtDNAcn that correlated with decreases in MYC protein in most cases. These findings provide evidence that, in late stage mCRPC, changes in mtDNAcn after drug treatment correlate with changes in MYC levels. This further supports the hypothesis that MYC activity may be a key factor controlling mtDNAcn in prostatic adenocarcinomas.

**Figure 6.**
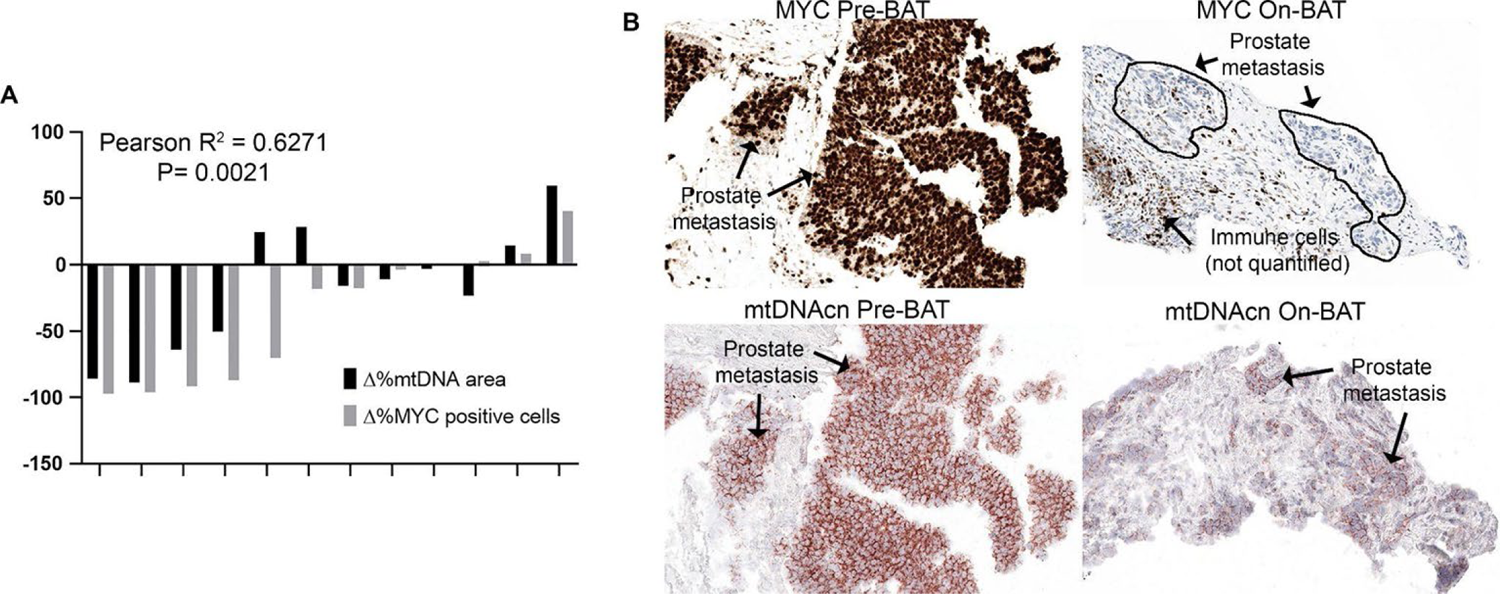
The decrease in MYC in response to SPA correlates with decreased mtDNAcn in mCRPC. (**A**) Each pair of bars shows the percentage change from the baseline biopsy in the percentage of MYC positive cells (by IHC staining for MYC protein) among all epithelial tumor cells by image analysis to the percentage change in mtDNA area by image analysis. There was a highly significant correlation (P=0.0021, R^2^=0.63). (**B**) Representative images of MYC-IHC and mtDNAcn-ISH for patients at the pretreatment time point and after 3 cycles of BAT. Notice that with both markers, image analysis was only performed in the epithelial compartments. MYC protein and mRNA quantification was reported previously for these cases (51). Original magnification, ∼x200.

### mtDNA replication as a potential downstream target of MYC in prostate cancer

The mechanisms by which MYC drives increased mtDNA levels in human and mouse PIN and human prostate cancer may be the result of increased mtDNA replication. In that case, inhibitors of mtDNA replication could provide a novel therapeutic approach for prostate cancer, similar to that being pursued currently in AML (36, 41).

To address whether MYC can drive mtDNA replication in prostate cancer cells (PC3), we inhibited MYC activity with a novel small molecule inhibitor MYCi975 (56, 62), followed by metabolic labeling with EdU for incorporation into newly synthesized DNA. Using an anti-5’-bromodeoxyuridine (BrdU) antibody that also binds to EdU (82) (**Fig. 7A**), we visualized the EdU signals by IHC. In untreated cells, we observed abundant punctate dot-like structures in the cytoplasm, consistent with mitochondrial localization. As a control for antibody specificity, these IHC signals were absent in cells not treated with EdU. In support of the hypothesis that these signals reflect mtDNA replication, the EdU-IHC signals were greatly reduced in cells treated with the mtDNA polymerase (POLG) inhibitor, ddC. Many cells also showed strong nuclear signals for EdU, indicating nuclear S phase labeling (**Fig. 7A**). After inhibition of MYC by MYCi975, there were reduced cytoplasmic signals, consistent with a decrease in mtDNA replication. There was also a decrease in nuclear DNA signals after MYC inhibition, which is expected since loss of MYC will also block nuclear DNA synthesis in prostate cancer cells (59). As a second approach, we performed qPCR for mtDNA and nuclear DNA (nDNA) after immunoprecipitation of EdU containing DNA. Using this assay, we also observed decreased EdU incorporation into mitochondrial genes when MYC levels were reduced (**Fig. 7B**). Taken together, the results of these two methods support the hypothesis that MYC activity controls mtDNA synthesis in prostate cancer cells.

**Figure 7.**
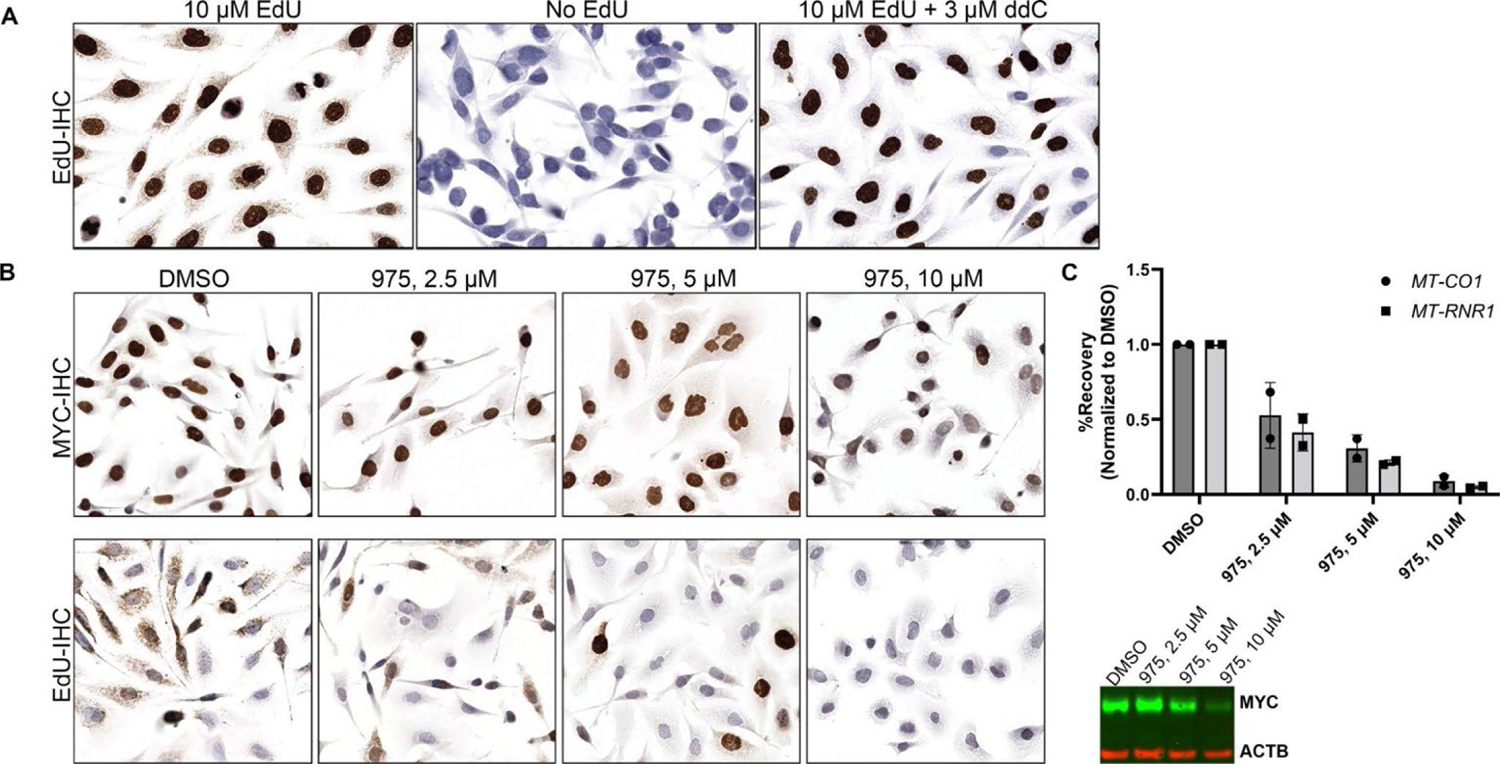
MYC activity controls mitochondrial DNA synthesis. (**A**) Representative images showing that the Anti-BrdU antibody staining is only present after PC3 cells are pretreated with EdU, demonstrating its specificity for EdU. Cells grown on chamber slides were metabolically labeled with EdU or no EdU and were then fixed and stained by IHC. Note the complete absence of staining in the middle panel in cells without EdU treatment and the marked reduction in cytoplasmic signals after treatment with the mtDNA replication inhibitor, ddC (right most panel). (**B-C**) When MYC activity and/or levels were decreased using MYCi975, decreased EdU incorporation into cytoplasmic or mitochondrial foci was observed by IHC (**B**) and separately by EdU immunoprecipitation followed by qPCR in PC3 cells (**C**). n = 2 independent biological replicates in IP. Representative Western blot results confirmed the decrease in MYC after drug treatment (**C**) in the samples subjected to IP. Original magnification for IHC images, x400 (**A**), x200 (**B**).

### mtDNA Replication Factors in Prostate Cancer

Genes required for mtDNA replication, which are all encoded by nuclear DNA, include *POLG* (the main DNA polymerase), *POLG2* (a processivity subunit), *TWNK* (the helicase referred to as Twinkle), *SSBP1* (the mtDNA specific single stranded binding protein), *TFAM* (encoding mitochondrial transcription factor A required for replication and transcription), *LIG3* (the DNA ligase), *PRIMPOL* (the DNA primase-polymerase), *TOP1MT* (encoding the mitochondrial topoisomerase) and *TOP3A* (a mitochondrial isoform encoding topoisomerase required to deconcatenate mtDNA circles). Germline mutations in the genes that encode these proteins lead to mtDNA depletion syndromes (83). A number of these genes have been previously shown to be MYC targets in other cell types (53, 84). To examine the potential regulation of these genes in prostate cancer by MYC, we interrogated mRNA levels of these genes after siRNA knockdown of MYC in 3 prostate cancer cell lines (59).

Most of the mtDNA replication genes were reduced after MYC knockdown (**Sup. Fig. 6A**). Publicly available ChIP-seq shows that MYC binding occurs within the promoter regions of multiple mtDNA replisome genes in a prostate cancer cell line; RNA-seq data showed their transcripts were reduced correspondingly (62) (**Sup. Fig. 6B**). Further, we examined the RNAseq data mentioned above from the U01 study. **Fig. 8** shows increased mRNA in primary prostate cancer lesions compared to matched benign for a number of genes encoding these mtDNA replication factors, especially Twinkle (*TWNK/C10ORF2*). Further, a number of these factors, including Twinkle, correlate with MYC expression in these primary tumors (**Fig. 8C**). When we examined RNAseq data from the COMBAT CRPC trial discussed above (51), a similar correlation with *MYC* and *TWNK* was present (**Sup Fig. 6C**). It is notable that this correlation between *MYC* and *TWNK* mRNA levels was also found in other cancer types seen in publically available data sets from the TCGA including those from human breast carcinoma, pancreatic adenocarcinoma and colorectal carcinoma (**Sup Fig. 7**). These findings suggest that a number of the necessary components of mtDNA replication machinery are positively regulated by MYC, are bound in their regulatory regions by MYC protein, are overexpressed in human prostatic cancer, and correlate with MYC levels in primary tumors and in mCRPC, providing further support that mtDNA replication may be an important downstream MYC target to investigate further.

**Figure 8.**
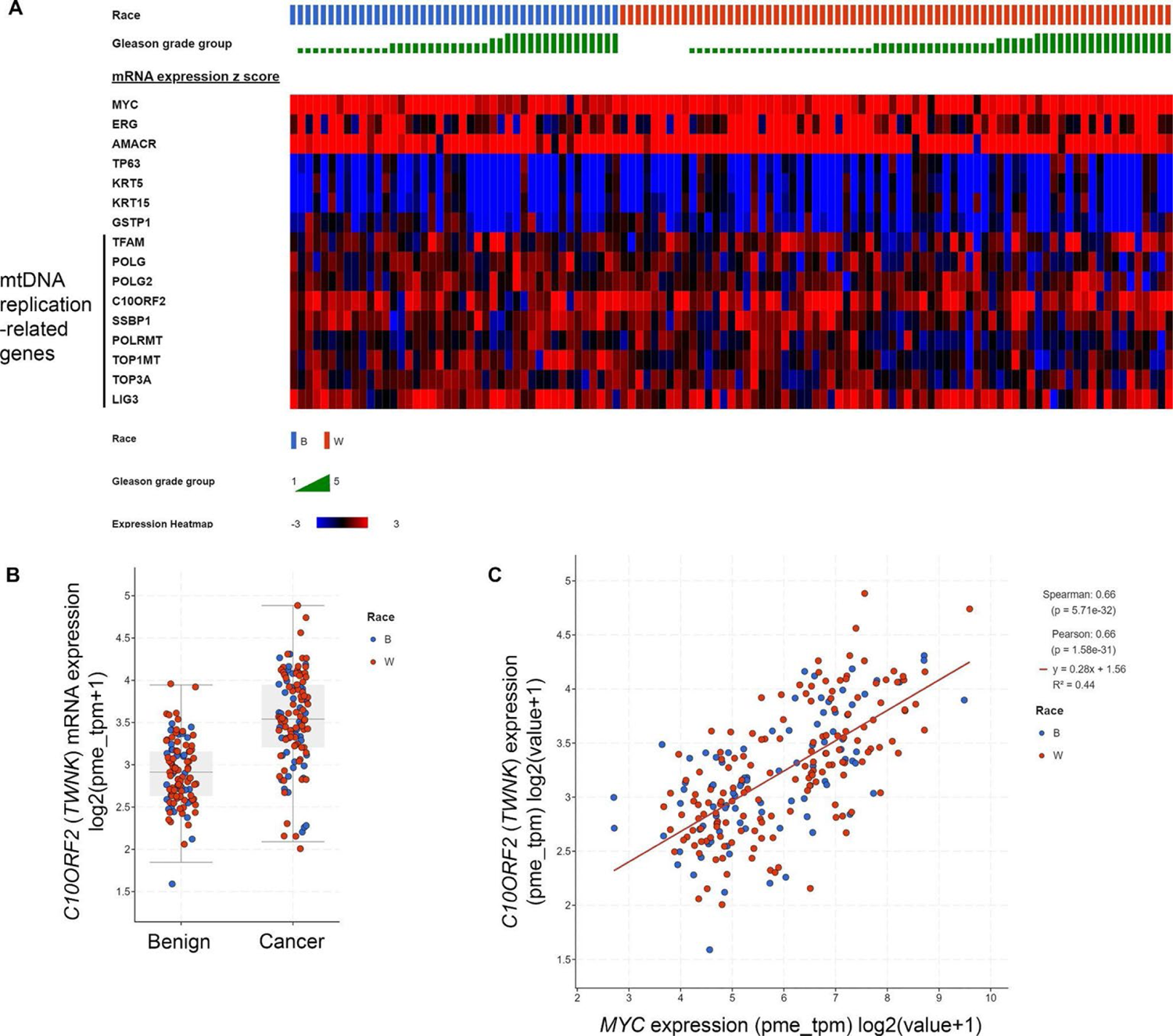
Genes encoding proteins required for mtDNA replication are upregulated in prostate cancer from human laser capture microdissected samples as determined by RNAseq. (**A**) Heatmap showing upregulation of transcripts in genes encoding mtDNA replication factors. Top line shows the Gleason grade group and the second line shows the patient race. *ERG*, *AMACR*, serve as controls for known upregulated genes in prostate cancer and *TP63*, *KRT5*, *KRT15* and *GSTP1* serve as controls for known downregulated (basal cell enriched) genes in prostate cancer. Genes directly involved in mtDNA replication are *TFAM*, *POLG* (mtDNA polymerase), *POLG2*, *C10ORF2* (*TWNK*), *SSBP1*, *LIG3*, *PRIMPOL* and *TOP3A*. (**B**) *TWNK/C10ORF2* mRNA was increased in cancer compared with benign (P<0.0001). (**C**) Correlation of *TWNK/C10ORF2* mRNA with *MYC* mRNA.

### Increased mtDNAcn was also seen in other cancer precursor lesions

To determine the applicability of our *in situ* approach and to ascertain how prevalent changes in mtDNAcn are in other cancer precursor lesions, we investigated mtDNAcn in precancerous lesions from the pancreas and large intestine. We applied the same CISH-IHC workflow on human pancreas tissues with pancreatic intraepithelial neoplasia (PanIN) and benign pancreatic ducts, as well as human colon tissues with tubular adenoma and normal colon mucosa. By visual inspection, PanIN lesions showed higher mtDNA *in situ* hybridization signals compared to their normal duct counterparts (**Figure 9A**). By image analysis, we observed approximately 3-fold higher %mtDNA area in PanIN compared to the normal ducts (**Figure 9B**). In the colonic samples, as recently reported (48), when present, the normal appearing colonic epithelium showed a marked gradation in mtDNAcn with cells located in the stem/proliferative compartments of crypts showing higher mtDNAcn compared to more differentiated cells towards the surface. In 7 of 8 adenomas, by visual inspection the mtDNA signals were more similar to, or apparently higher than, the levels observed in the normal appearing stem cell compartment (**Figure 9C**). These strong mtDNA signals occurred even towards the surface of the adenomatous epithelium in a manner reminiscent of topographical infidelity of proliferation markers and other proteins in colorectal adenomas, including MYC protein expression (**Figure 9D**) (85, 86). Overall, these results suggest that increased mtDNAcn may accompany precancerous neoplastic lesion development across multiple organ systems.

**Figure 9.**
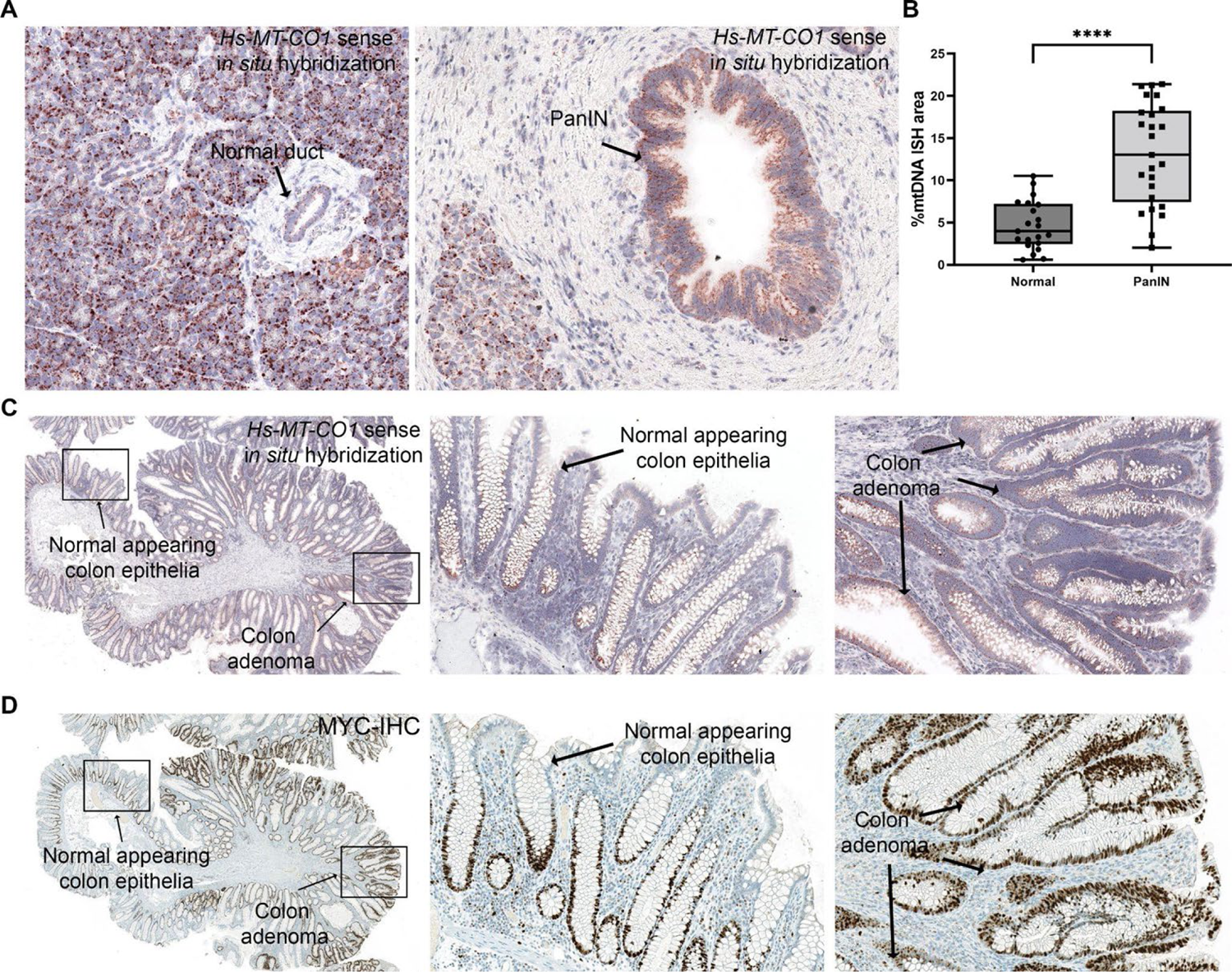
Increased mtDNAcn is commonly found in other cancer precursor lesions. (**A**) Representative *in situ* hybridization results of a normal pancreatic duct versus a PanIN. (**B**) Quantification of mtDNAcn in normal ducts and PanINs using image analysis. Each symbol represents a region of the tissue type specified. (**C-D**) Representative images of mtDNA *in situ* hybridization and MYC IHC in a colon sample containing an adenomatous polyp. Boxed areas in the left-most images are shown as higher magnification in the middle and right panels. Original magnification, x200 (**A**), x20 (**C**-**D**), x400 (middle and right panels in **C** and **D**). ****P < 0.0001.

## Discussion

To overcome the limitations inherent in most mtDNAcn measurements in tumor/normal pairs, we developed a multiplex chromogenic *in situ* hybridization and IHC approach to quantify mtDNAcn in specific cell populations. This assay uncovered consistent increases in mtDNAcn in high grade PIN. In addition, we found increases and marked intratumoral heterogeneity in mtDNAcn in most primary prostatic adenocarcinomas and further increases in advanced mCRPC. The increased mtDNAcn in primary prostate cancer was independently verified using whole genome sequencing (WGS) of DNA isolated after LCM, as well as by quantitative real-time PCR. Since three separate complementary approaches produced highly similar results, our findings should resolve previous contradictory publications regarding differences in mtDNAcn in prostatic cancers as compared with matched benign normal appearing prostate epithelium. The increased mtDNAcn in prostate cancer cells is functionally active, as shown by corresponding increases in mtRNAs and mitochondrial electron transport chain activity, observed by *in situ* enzymatic activity. Mechanistically, increased mtDNAcn in mouse prostatic luminal epithelial cells *in vivo* is driven by forced overexpression of MYC. Additionally, reductions in mtDNAcn correlate with reductions in MYC that are observed in mCRPC clinical samples after treatment with supraphysiological testosterone (51). Since MYC is known to be consistently overexpressed in human PIN (compared with normal prostate luminal cells) and most primary and metastatic castration resistant prostatic adenocarcinomas (51, 52, 66, 73), these findings implicate MYC overexpression as causing mtDNAcn increases in these lesions. We also report that a key mechanism by which MYC drives increased mtDNAcn is through increased mtDNA replication, potentially via positive regulation of mtDNA replication factors, including the helicase, TWINKLE. Finally, our *in situ* approach also identified increases in mtDNAcn in neoplastic precursor lesions in the colon (adenomas) and pancreas (PanINs), demonstrating the broad applicability of this method.

### Comparisons with other prostate cancer studies

Our findings are consistent with the findings of Mizumachi et al. (43) Our *in situ* method and their single cell LCM approach both measure mtDNAcn with single cell resolution. Our findings also generally agree with a more recent study using whole genome sequencing data from TCGA samples that showed a trend for increased mtDNAcn in prostate cancer versus benign matched tissues (45). Although Reznik et al. reported that there was no correlation between mtDNAcn and mtRNA expression in prostate cancer using sequencing methods (87), we observed that at least for the 3 mtRNAs we examined in this study, their spatial patterns and expression levels highly correlate with those of mtDNAcn. The present findings differ with those of Schöpf et al., (18) who found no overall increase in mtDNAcn in cancer vs. normal in the prostate, although they did report an increase in mtDNAcn in higher grade tumors as compared with the lower grade ones. By contrast, we found consistently increased mtDNAcn across all grade groups. Nevertheless, in our study, the Gleason grade of all of the cases with the highest levels of mtDNA in primary tumors were greater than Grade Group 1, and we found a further increase in metastatic castration resistant lesions (**Fig. 2** and **Fig. 3**). Schöpf et al. (18) used tissue punch biopsies to isolate tumor and benign tissues, which provides a lower level of tumor purity than our laser capture based method used for our WGS approach, and this could help explain the differences. Our findings differed from those of Koochekpour et al. (44), since we found no difference in Black vs. White men in tumor mtDNAcn, while their study did.

### Basal vs. luminal cells

In the present study we found that basal cells have approximately 5-fold higher %mtDNA area as compared with normal luminal cells. In comparing mtDNA cellular levels in HGPIN and invasive adenocarcinomas to normal, there was a clear increase overall, even when including normal basal cells in the normal regions (**Fig 2**). Although some prostatic adenocarcinomas appear to possess gene expression signatures, or modules, that resemble basal cells to a degree (88–90), most evidence suggests that the cell of origin for prostate carcinoma is a luminal cell (66, 91–93). When compared specifically to benign normal appearing luminal cells, there was a more striking increase in HGPIN luminal cells (**Sup. Fig. 8**).

### MYC may drive mtDNA replication

Prior studies have shown that MYC can regulate mitochondrial biogenesis and mitochondrial DNA synthesis (53, 84). Wang et al., reported that during *Drosophila* oogenesis, *myc* regulated mtDNA replication and COX enzyme activity, while a hypomorphic *myc* allele led to reduced levels of both (84). They further suggested that such changes may occur through myc’s role as a master transcription factor, since a majority of the genes encoding proteins of the electron transport chain, mtDNA replication and transcription were downregulated in flies with the hypomorphic *myc* alleles. Our results mirrored theirs as we observed similar trends when examining microarray data after knockdown of MYC in human prostate cancer cell lines, as well as RNAseq results from a clinical trial where responding patients showed reduced MYC was accompanied reduced mRNA levels in genes such as *TWNK* (**Supplemental Fig. 6**). Also, a number of mtDNA replication-related genes are upregulated in human prostate cancer and *TWNK* levels correlate with *MYC* mRNA (**Fig. 8** and **Sup. Fig. 6C**). Taken together, these findings suggest an essential role of MYC in mtDNA replication and mitochondrial function in both *Drosophila* normal ovarian tissues and prostate cancer development and progression. Further study is required to assess whether increased mtDNAcn (or de novo mtDNA synthesis) is required for tumor development and progression in MYC-driven cancers.

### Why is mtDNAcn increased?

The overall biological significance behind the observed increase of mtDNAcn in HGPIN, prostate cancer, and in mCRPC, is currently unknown. We hypothesize that increased mtDNAcn is necessary in the process of carcinogenesis (40, 41). For instance, we hypothesize that higher mtDNAcn drives mitochondrial biogenesis to increase mitochondrial activity needed for the increase in macromolecular synthesis required for cell growth and replication. Since our mCRPC samples generally showed increased mtDNAcn compared to primary prostate cancer, we hypothesize that the cells with high mtDNAcn have a growth advantage and thus may be selected during disease progression. In support of this, we found reduced mtDNAcn in matched samples from patients before and after SPA that correlated with clinical response, Ki67 levels and MYC levels. Studies over several decades have indicated that, compared with normal prostate, prostate cancer cells show increased mitochondrial functions that are important for cell proliferation, and our overall results are concordant with these observations (94–98).

An alternative hypothesis is that, since mtDNA mutations are common in prostate cancer (10, 15, 18, 87), some of which may be deleterious to mitochondrial functions, increased mtDNAcn may reflect compensatory upregulation of mtDNA levels to help maintain these compromised mitochondrial functions. There is a potential precedent for this in prostate cancer as seen by Schöpf et al., in which they found higher mtDNAcn in cases showing potentially deleterious mtDNA mutations (18). In that study, short term cultures of fresh prostate cancer tissue showed an altered mitochondrial oxidative phosphorylation capacity including a decrease in N-pathway (NADH through Complex I, III, IV) capacity, and an increase in S-pathways (succinate from complex II-IV) capacity, which was accentuated in cases with higher levels of heteroplasmy of potentially deleterious mtDNA mutations in proteins encoding complex I components (18). Furthermore, mtDNA mutations, or nuclear-encoded genes that can alter the TCA cycle flux, can result in increases in “oncometabolites’’ that can drive epigenetic changes such as DNA CpG methylation that can provide a selective growth advantage to the neoplastic cells (25), despite having somewhat compromised overall mitochondrial function. Thus, in this case, there could also be compensatory increases in mtDNAcn to maintain mitochondrial function.

### Translational Relevance and Future Directions

The discovery of functional mitochondrial reprogramming geared for biomass generation and proliferation in many cancers (1) has nominated pharmacological blockade of mitochondrial oxidative phosphorylation, mtDNA replication, and mitochondrial translation (28, 99) as promising and novel anti-cancer therapeutic avenues. Based upon the present findings, we hypothesize that developing novel drugs to inhibit required mtDNA replication machinery components, including the helicase Twinkle (encoded by *TWNK*), SSBP1 and mitochondrial topoisomerases, may represent new therapeutic approaches for cancer treatment and/or prevention. This concept is analogous to the recent credentialing of POLMRT, the mtRNA polymerase, as a novel cancer drug target (100). Related to this, it has been hypothesized that those tumors with high levels of activity of canonical mitochondrial functions, such as TCA cycle and oxidative phosphorylation, may harbor dependencies that could be most susceptible to mitochondrial inhibition therapies (27, 37, 101). Since mtDNAcn regulation is associated with controlling mitochondrial biogenesis and function, accurate measurements of mtDNAcn in tumors and their precursors using our *in situ* approach may help determine which cancers are most appropriate for mitochondrial therapies. Also, the fact that we observed elevated mtDNAcn in precursor lesions from 3 different tumor types, and others have seen in head and neck cancers (49), suggest that blockade of mitochondrial activities may be an important new approach for precision prevention and cancer interception strategies. Future methods that can measure mtDNA using single cell based sequencing methods may also have an important role here (102), although they would lose critical spatial information that is a key feature in our approach. The method developed in this study is straightforward and should be readily portable to automated *in situ* assays combined with IHC, when appropriate, that could be implemented routinely in research laboratories and clinical pathology laboratories if additional future studies support clinical utility.

## Author contributions

JC and AMD conceptualized and designed the project. JC, QZ, JBV, JLH, LT, BO, TJ, AV, TCL, IK, RW, MM, SRD, KP, SW, RH, EA, AG,SY and AMD acquired, analyzed and interpreted data. JC and AMD wrote the original draft of the manuscript. All authors were responsible for reviewing and editing the manuscript.

## Acknowledgements

We thank Hiromi Sesaki for providing the detailed cytochrome c oxidase and succinate dehydrogenase tissue based enzyme activity assay protocol. We thank Sarah Wheelan for helping with WGS and RNAseq analysis. We thank Alan Meeker for helping with fluorescent whole slide scanning. This work is supported by NIH/National Cancer Institute (NCI) Specialized Programs of Research Excellence (SPORE) in Prostate Cancer grant P50CA58236 (AMD), NIH/NCI grant U01 CA196390 (AMD, SY, KP.), NIH/NCI U54 CA274370 (AMD, SY, KP.), U.S. Department of Defense Prostate Cancer Research Program (PCRP) grant W81XWH-18-2-0015 (AMD); The Johns Hopkins Sidney Kimmel Comprehensive Cancer Center Oncology Tissue Services Laboratory is supported by NIH/NCI grant P30 CA006973; The Patrick C. Walsh Prostate Cancer Research Fund at Johns Hopkins (AMD, SY).

## References

1. Vander Heiden MG, Cantley LC, Thompson CB. Understanding the Warburg effect: the metabolic requirements of cell proliferation. Science 2009;324(5930):1029–1033.

2. Wallace DC. Mitochondria and cancer. Nat. Rev. Cancer 2012;12(10):685–698.

3. López-Otín C, et al. The hallmarks of aging. Cell 2013;153(6):1194–1217.

4. O’Hara R, et al. Quantitative mitochondrial DNA copy number determination using droplet digital PCR with single cell resolution. Genome Res. [published online ahead of print: September 23, 2019]; doi:10.1101/gr.250480.119

5. Jones JB, et al. Detection of mitochondrial DNA mutations in pancreatic cancer offers a “mass”-ive advantage over detection of nuclear DNA mutations. Cancer Res. 2001;61(4):1299–1304.

6. Eng C, et al. A role for mitochondrial enzymes in inherited neoplasia and beyond. Nat. Rev. Cancer 2003;3(3):193–202.

7. Neumann HPH, et al. Distinct clinical features of paraganglioma syndromes associated with SDHB and SDHD gene mutations. JAMA 2004;292(8):943–951.

8. Pasini B, et al. Clinical and molecular genetics of patients with the Carney-Stratakis syndrome and germline mutations of the genes coding for the succinate dehydrogenase subunits SDHB, SDHC, and SDHD. Eur. J. Hum. Genet. 2008;16(1):79–88.

9. Crooks DR, et al. Mitochondrial DNA alterations underlie an irreversible shift to aerobic glycolysis in fumarate hydratase-deficient renal cancer. Sci. Signal. 2021;14(664):eabc4436.

10. Petros JA, et al. mtDNA mutations increase tumorigenicity in prostate cancer. Proc. Natl. Acad. Sci. U. S. A. 2005;102(3):719–724.

11. Arnold RS, et al. Mitochondrial DNA mutation stimulates prostate cancer growth in bone stromal environment. Prostate 2009;69(1):1–11.

12. Kloss-Brandstätter A, et al. Somatic Mutations throughout the Entire Mitochondrial Genome Are Associated with Elevated PSA Levels in Prostate Cancer Patients. The American Journal of Human Genetics 2010;87(6):802–812.

13. Arnold RS, et al. An inherited heteroplasmic mutation in mitochondrial gene COI in a patient with prostate cancer alters reactive oxygen, reactive nitrogen and proliferation. Biomed Res. Int. 2013;2013:239257.

14. McCrow JP, et al. Spectrum of mitochondrial genomic variation and associated clinical presentation of prostate cancer in South African men. Prostate 2016;76(4):349–358.

15. Hopkins JF, et al. Mitochondrial mutations drive prostate cancer aggression. Nat. Commun. 2017;8(1):656.

16. Kalsbeek AMF, et al. Mitochondrial genome variation and prostate cancer: a review of the mutational landscape and application to clinical management. Oncotarget 2017;8(41):71342–71357.

17. Philley JV, et al. Complex-I alteration and enhanced mitochondrial fusion are associated with prostate cancer progression. J. Cell. Physiol. 2016;231(6):1364–1374.

18. Schöpf B, et al. OXPHOS remodeling in high-grade prostate cancer involves mtDNA mutations and increased succinate oxidation. Nat. Commun. 2020;11(1):1487.

19. Maki J, et al. Mitochondrial genome deletion aids in the identification of false- and true-negative prostate needle core biopsy specimens. Am. J. Clin. Pathol. 2008;129(1):57–66.

20. Zheng J. Energy metabolism of cancer: Glycolysis versus oxidative phosphorylation (Review). Oncol. Lett. 2012;4(6):1151–1157.

21. Luengo A, et al. Increased demand for NAD+ relative to ATP drives aerobic glycolysis. Mol. Cell 2021;81(4):691–707.e6.

22. Lunt SY, Vander Heiden MG. Aerobic glycolysis: meeting the metabolic requirements of cell proliferation. Annu Rev Cell Dev Biol 2011;27:441–464.

23. Koppenol WH, Bounds PL, Dang CV. Otto Warburg’s contributions to current concepts of cancer metabolism. Nat Rev Cancer 2011;11(5):325–337.

24. Pavlova NN, Thompson CB. The Emerging Hallmarks of Cancer Metabolism. Cell Metab. 2016;23(1):27– 47.

25. Zong W-X, Rabinowitz JD, White E. Mitochondria and Cancer. Mol. Cell 2016;61(5):667–676.

26. Vander Heiden MG, DeBerardinis RJ. Understanding the Intersections between Metabolism and Cancer Biology. Cell 2017;168(4):657–669.

27. Ward PS, Thompson CB. Metabolic reprogramming: a cancer hallmark even warburg did not anticipate. Cancer Cell 2012;21(3):297–308.

28. Vasan K, Werner M, Chandel NS. Mitochondrial Metabolism as a Target for Cancer Therapy. Cell Metab. [published online ahead of print: July 8, 2020]; doi:10.1016/j.cmet.2020.06.019

29. Weinberg F, et al. Mitochondrial metabolism and ROS generation are essential for Kras-mediated tumorigenicity. Proc Natl Acad Sci U S A 2010;107(19):8788–8793.

30. Sullivan LB, et al. Supporting aspartate biosynthesis is an essential function of respiration in proliferating cells. Cell 2015;162(3):552–563.

31. Zhu J, Thompson CB. Metabolic regulation of cell growth and proliferation. Nat. Rev. Mol. Cell Biol. 2019;20(7):436–450.

32. Bajzikova M, et al. Reactivation of Dihydroorotate Dehydrogenase-Driven Pyrimidine Biosynthesis Restores Tumor Growth of Respiration-Deficient Cancer Cells. Cell Metab 2019;29(2):399–416.e10.

33. Pavlova NN, Zhu J, Thompson CB. The hallmarks of cancer metabolism: Still emerging. Cell Metab. 2022;34(3):355–377.

34. Clay Montier LL, Deng JJ, Bai Y. Number matters: control of mammalian mitochondrial DNA copy number. J. Genet. Genomics 2009;36(3):125–131.

35. Castellani CA, et al. Thinking outside the nucleus: Mitochondrial DNA copy number in health and disease. Mitochondrion 2020;53:214–223.

36. Boultwood J, et al. Amplification of mitochondrial DNA in acute myeloid leukaemia. Br. J. Haematol. 1996;95(2):426–431.

37. Skrtić M, et al. Inhibition of mitochondrial translation as a therapeutic strategy for human acute myeloid leukemia. Cancer Cell 2011;20(5):674–688.

38. Cormio A, et al. Mitochondrial DNA content and mass increase in progression from normal to hyperplastic to cancer endometrium. BMC Res. Notes 2012;5:279.

39. Lin C-S, et al. The role of mitochondrial DNA alterations in esophageal squamous cell carcinomas. J. Thorac. Cardiovasc. Surg. 2010;139(1):189–197.e4.

40. Sun X, et al. Increased mtDNA copy number promotes cancer progression by enhancing mitochondrial oxidative phosphorylation in microsatellite-stable colorectal cancer. Signal Transduct Target Ther 2018;3:8.

41. Liyanage SU, et al. Leveraging increased cytoplasmic nucleoside kinase activity to target mtDNA and oxidative phosphorylation in AML. Blood 2017;129(19):2657–2666.

42. Yap TA, et al. Complex I inhibitor of oxidative phosphorylation in advanced solid tumors and acute myeloid leukemia: phase I trials. Nat. Med. 2023;29(1):115–126.

43. Mizumachi T, et al. Increased distributional variance of mitochondrial DNA content associated with prostate cancer cells as compared with normal prostate cells. Prostate 2008;68(4):408–417.

44. Koochekpour S, et al. Reduced mitochondrial DNA content associates with poor prognosis of prostate cancer in African American men. PLoS One 2013;8(9):e74688.

45. Reznik E, et al. Mitochondrial DNA copy number variation across human cancers. Elife 2016;5. doi:10.7554/eLife.10769

46. Higuchi M, et al. Mitochondrial DNA determines androgen dependence in prostate cancer cell lines. Oncogene 2006;25(10):1437–1445.

47. Yuan Y, et al. Comprehensive molecular characterization of mitochondrial genomes in human cancers. Nat. Genet. [published online ahead of print: February 5, 2020]; doi:10.1038/s41588-019-0557-x

48. Chen J, et al. An in Situ Atlas of Mitochondrial DNA in Mammalian Tissues Reveals High Content in Stem/Progenitor Cells. Am. J. Pathol. [published online ahead of print: April 15, 2020]; doi:10.1016/j.ajpath.2020.03.018

49. Kim MM, et al. Mitochondrial DNA quantity increases with histopathologic grade in premalignant and malignant head and neck lesions. Clin. Cancer Res. 2004;10(24):8512–8515.

50. Abate-Shen C, Shen MM. Molecular genetics of prostate cancer. Genes Dev. 2000;14(19):2410–2434.

51. Sena LA, et al. Prostate cancer androgen receptor activity dictates efficacy of bipolar androgen therapy through MYC. J. Clin. Invest. [published online ahead of print: October 4, 2022]; doi:10.1172/JCI162396

52. Guo H, et al. Androgen receptor and MYC equilibration centralizes on developmental super-enhancer. Nat. Commun. 2021;12(1):1–18.

53. Li F, et al. Myc stimulates nuclearly encoded mitochondrial genes and mitochondrial biogenesis. Mol. Cell. Biol. 2005;25(14):6225–6234.

54. Ellwood-Yen K, et al. Myc-driven murine prostate cancer shares molecular features with human prostate tumors. Cancer Cell 2003;4(3):223–238.

55. Ozbek B, et al. Multiplex immunohistochemical phenotyping of T cells in primary prostate cancer. Prostate [published online ahead of print: February 21, 2022]; doi:10.1002/pros.24315

56. Han H, et al. Small-Molecule MYC Inhibitors Suppress Tumor Growth and Enhance Immunotherapy. Cancer Cell [published online ahead of print: October 22, 2019]; doi:10.1016/j.ccell.2019.10.001

57. Jiang M, et al. The mitochondrial single-stranded DNA binding protein is essential for initiation of mtDNA replication. Sci Adv 2021;7(27). doi:10.1126/sciadv.abf8631

58. Baena-Del Valle JA, et al. MYC drives overexpression of telomerase RNA (hTR/TERC) in prostate cancer. J. Pathol. 2018;244(1):11–24.

59. Koh CM, et al. Alterations in nucleolar structure and gene expression programs in prostatic neoplasia are driven by the MYC oncogene. Am. J. Pathol. 2011;178(4):1824–1834.

60. Korotkevich G, et al. Fast gene set enrichment analysis. bioRxiv 2021;060012.

61. Rath S, et al. MitoCarta3.0: an updated mitochondrial proteome now with sub-organelle localization and pathway annotations. Nucleic Acids Res. 2021;49(D1):D1541–D1547.

62. Holmes AG, et al. A MYC inhibitor selectively alters the MYC and MAX cistromes and modulates the epigenomic landscape to regulate target gene expression. Sci Adv 2022;8(17):eabh3635.

63. Gao J, et al. Integrative analysis of complex cancer genomics and clinical profiles using the cBioPortal. Sci. Signal. 2013;6(269):l1.

64. Cerami E, et al. The cBio cancer genomics portal: an open platform for exploring multidimensional cancer genomics data. Cancer Discov. 2012;2(5):401–404.

65. David G. Bostwick, Liang Cheng, Isabelle Meiers. Neoplasms of the Prostate. In: David G. Bostwick LC ed. Urologic Surgical Pathology, Third Edition. Elsevier Saunders; 2014:410–531

66. Trabzonlu L, et al. Molecular Pathology of High-Grade Prostatic Intraepithelial Neoplasia: Challenges and Opportunities. Cold Spring Harb. Perspect. Med. [published online ahead of print: August 6, 2018]; doi:10.1101/cshperspect.a030403

67. Haffner MC, et al. Molecular evidence that invasive adenocarcinoma can mimic prostatic intraepithelial neoplasia (PIN) and intraductal carcinoma through retrograde glandular colonization. J. Pathol. 2016;238(1):31–41.

68. Gumuskaya B, et al. Assessing the order of critical alterations in prostate cancer development and progression by IHC: further evidence that PTEN loss occurs subsequent to ERG gene fusion. Prostate Cancer Prostatic Dis. 2013;16(2):209–215.

69. Maughan BL, et al. Pharmacodynamic study of the oral hedgehog pathway inhibitor, vismodegib, in patients with metastatic castration-resistant prostate cancer. Cancer Chemother. Pharmacol. 2016;78(6):1297–1304.

70. Morrish F, Hockenbery D. MYC and mitochondrial biogenesis. Cold Spring Harb. Perspect. Med. 2014;4(5). doi:10.1101/cshperspect.a014225

71. Gurel B, et al. Nuclear MYC protein overexpression is an early alteration in human prostate carcinogenesis. Mod. Pathol. 2008;21(9):1156–1167.

72. Koh CM, et al. MYC and Prostate Cancer. Genes Cancer 2010;1(6):617–628.

73. Arriaga JM, et al. A MYC and RAS co-activation signature in localized prostate cancer drives bone metastasis and castration resistance. Nature Cancer [published online ahead of print: October 19, 2020]; doi:10.1038/s43018-020-00125-0

74. Iwata T, et al. MYC overexpression induces prostatic intraepithelial neoplasia and loss of Nkx3.1 in mouse luminal epithelial cells. PLoS One 2010;5(2):e9427.

75. Hubbard GK, et al. Combined MYC Activation and Pten Loss Are Sufficient to Create Genomic Instability and Lethal Metastatic Prostate Cancer. Cancer Res. 2016;76(2):283–292.

76. Schweizer MT, et al. Effect of bipolar androgen therapy for asymptomatic men with castration-resistant prostate cancer: results from a pilot clinical study. Sci. Transl. Med. 2015;7(269):269ra2.

77. Teply BA, et al. Bipolar androgen therapy in men with metastatic castration-resistant prostate cancer after progression on enzalutamide: an open-label, phase 2, multicohort study. Lancet Oncol. 2018;19(1):76–86.

78. Markowski, et al. A multicohort open-label phase II trial of bipolar androgen therapy in men with metastatic castration-resistant prostate cancer (RESTORE): a comparison of …. Eur. Urol. https://www.sciencedirect.com/science/article/pii/S0302283820304711. cited

79. Sena LA, et al. Bipolar androgen therapy sensitizes castration-resistant prostate cancer to subsequent androgen receptor ablative therapy. Eur. J. Cancer 2021;144:302–309.

80. Denmeade SR, et al. TRANSFORMER: A Randomized Phase II Study Comparing Bipolar Androgen Therapy Versus Enzalutamide in Asymptomatic Men With Castration-Resistant Metastatic Prostate Cancer. J. Clin. Oncol. 2021;39(12):1371–1382.

81. Markowski MC, et al. Overall survival (OS) and biomarker results from combat: A phase 2 study of bipolar androgen therapy (BAT) plus nivolumab for patients with metastatic castrate-resistant prostate cancer (mCRPC). J. Clin. Oncol. 2022;40(16_suppl):5064–5064.

82. Liboska R, et al. Most anti-BrdU antibodies react with 2’-deoxy-5-ethynyluridine -- the method for the effective suppression of this cross-reactivity. PLoS One 2012;7(12):e51679.

83. McKinney EA, Oliveira MT. Replicating animal mitochondrial DNA. Genet. Mol. Biol. 2013;36(3):308–315.

84. Wang Z-H, et al. Electron transport chain biogenesis activated by a JNK-insulin-Myc relay primes mitochondrial inheritance in Drosophila. Elife 2019;8:e49309.

85. Polyak K, et al. Early Alteration of Cell-Cycle-Regulated Gene. Am J Pathol 1996;149:381–387.

86. De Marzo AM, et al. Abnormal regulation of DNA methyltransferase expression during colorectal carcinogenesis. Cancer Res. 1999;59(16):3855–3860.

87. Reznik E, et al. Mitochondrial respiratory gene expression is suppressed in many cancers. Elife 2017;6. doi:10.7554/eLife.21592

88. Goldstein AS, et al. Identification of a cell of origin for human prostate cancer. Science 2010;329(5991):568–571.

89. Smith BA, et al. A basal stem cell signature identifies aggressive prostate cancer phenotypes. Proc. Natl. Acad. Sci. U. S. A. 2015;112(47):E6544–52.

90. Aggarwal R, et al. Prognosis Associated With Luminal and Basal Subtypes of Metastatic Prostate Cancer. JAMA Oncol [published online ahead of print: September 23, 2021]; doi:10.1001/jamaoncol.2021.3987

91. Wang ZA, et al. Luminal cells are favored as the cell of origin for prostate cancer. Cell Rep. 2014;8(5):1339–1346.

92. De Marzo AM, et al. Premalignancy in Prostate Cancer: Rethinking What we Know. Cancer Prev. Res. 2016;9(8):648–656.

93. Zhang D, et al. Prostate Luminal Progenitor Cells in Development and Cancer. Trends Cancer Res. 2018;4(11):769–783.

94. Khan MA, et al. Growth pattern and citrate production in organ cultures of adult rat ventral prostate. Prostate 1982;3(4):391–403.

95. Grupp K, et al. High mitochondria content is associated with prostate cancer disease progression. Mol. Cancer 2013;12(1):145.

96. Dakubo GD, et al. Altered metabolism and mitochondrial genome in prostate cancer. J. Clin. Pathol. 2006;59(1):10–16.

97. Bader DA, et al. Mitochondrial pyruvate import is a metabolic vulnerability in androgen receptor-driven prostate cancer. Nat Metab 2019;1(1):70–85.

98. Bader DA, McGuire SE. Tumour metabolism and its unique properties in prostate adenocarcinoma. Nat. Rev. Urol. [published online ahead of print: February 28, 2020]; doi:10.1038/s41585-020-0288-x

99. Delaunay S, et al. Mitochondrial RNA modifications shape metabolic plasticity in metastasis | Nature. *Nature* 2022;607(7919):593–603.

100. Bonekamp NA, et al. Small-molecule inhibitors of human mitochondrial DNA transcription. Nature 2020;1–5.

101. Masoud R, et al. Targeting Mitochondrial Complex I Overcomes Chemoresistance in High OXPHOS Pancreatic Cancer. Cell Rep Med 2020;1(8):100143.

102. Lareau CA, et al. Massively parallel single-cell mitochondrial DNA genotyping and chromatin profiling. Nat. Biotechnol. 2021;39(4):451–461.

