## Supplemental Figures and Tables for "MYC-driven increases in mitochondrial DNA copy number occur early and persist throughout prostatic cancer progression"

### Supplemental Figures and Supplemental Figure Legends

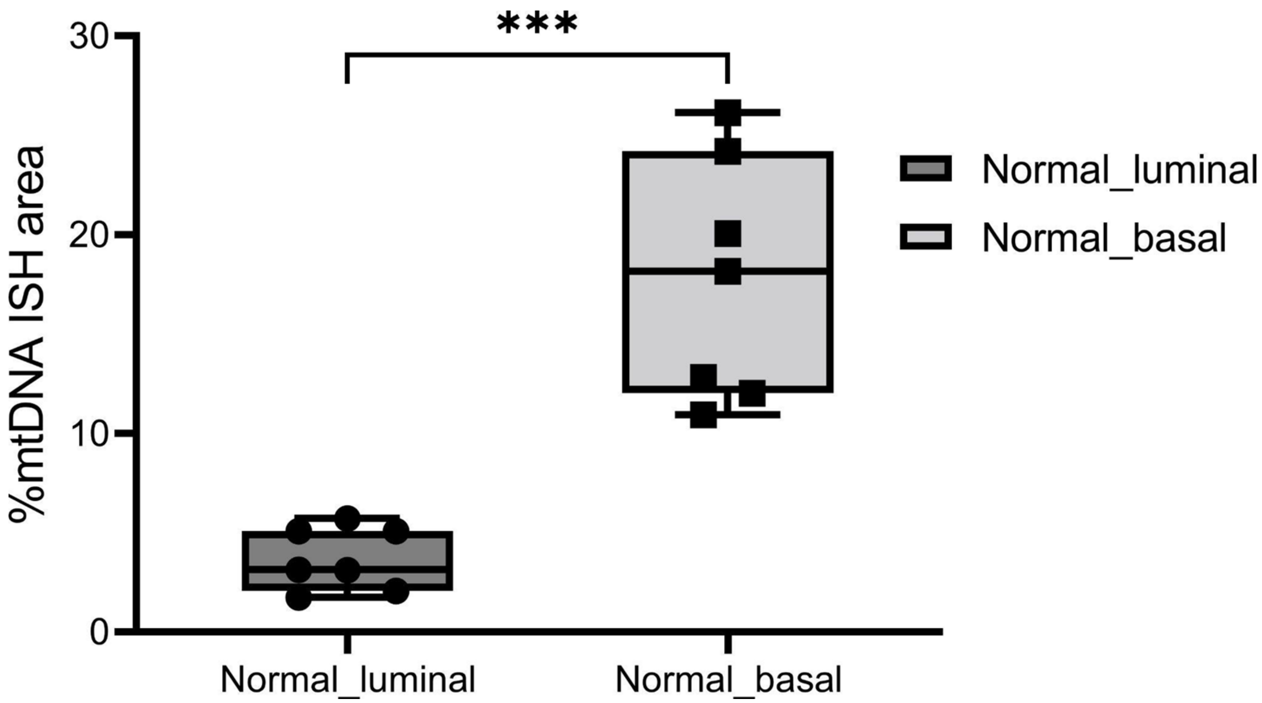


**Supplemental Figure 1. Image analysis results showing higher mtDNAcn in normal basal cells compared to normal luminal cells.** The center line in the box shows the median %mtDNA area in each group. n = 7 regions from 3 patients for each cell type. ***P < 0.0002.

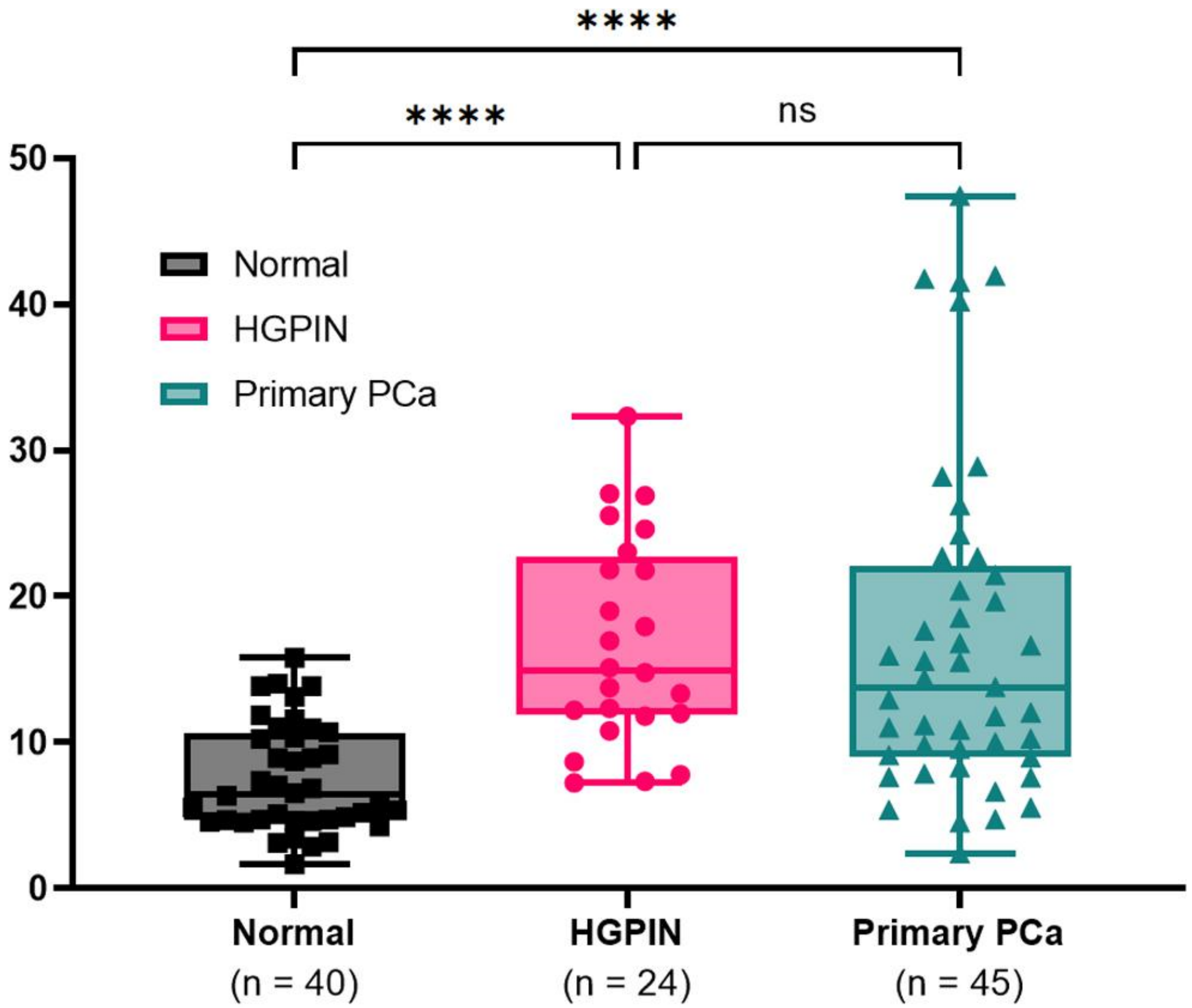


**Supplemental Figure 2**. **Heterogeneity of mtDNAcn in human prostate.** Quantitative image analysis results from individual regions of normal, HGPIN and primary prostate carcinoma. Each symbol represents an individual region of interest examined. ****P < 0.0001.

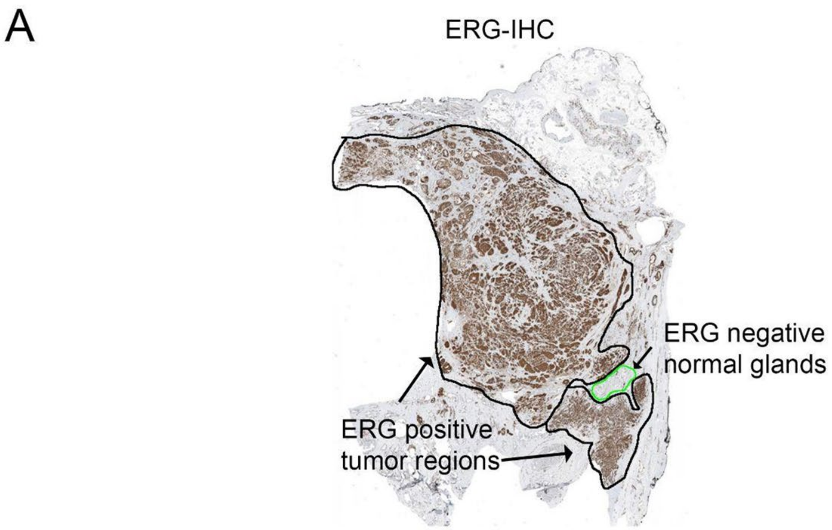


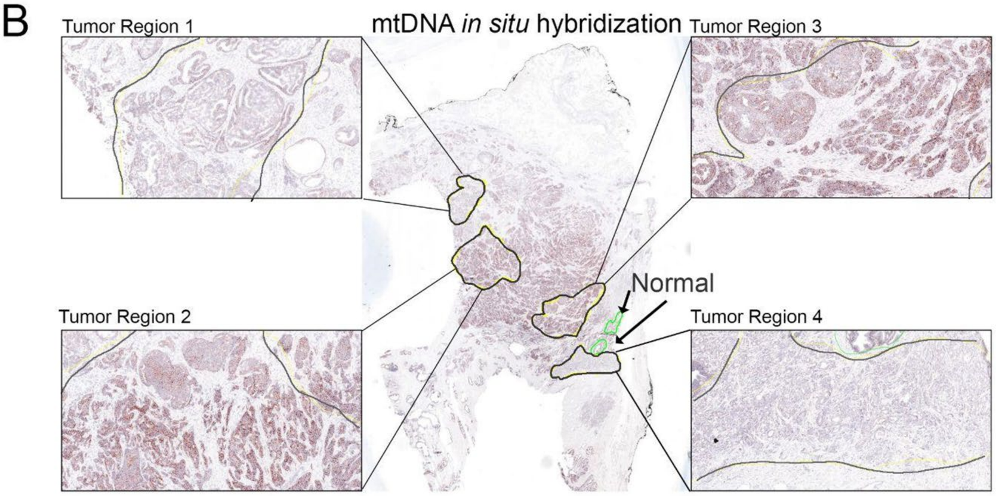


**Supplemental Figure 3. Intratumoral heterogeneity in a single prostatic tumor nodule.** (**A**) Low power view of a prostatectomy specimen with tumor nodule that is homogeneously positive for ERG protein, indicative of an *ERG* gene fusion. Original magnification, x10. (**B**) Marked heterogeneity of mtDNA *in situ* hybridization signals from different subregions of this tumor using an adjacent slide to that shown in **A**. Original magnification, x10 and x70 (boxed regions).

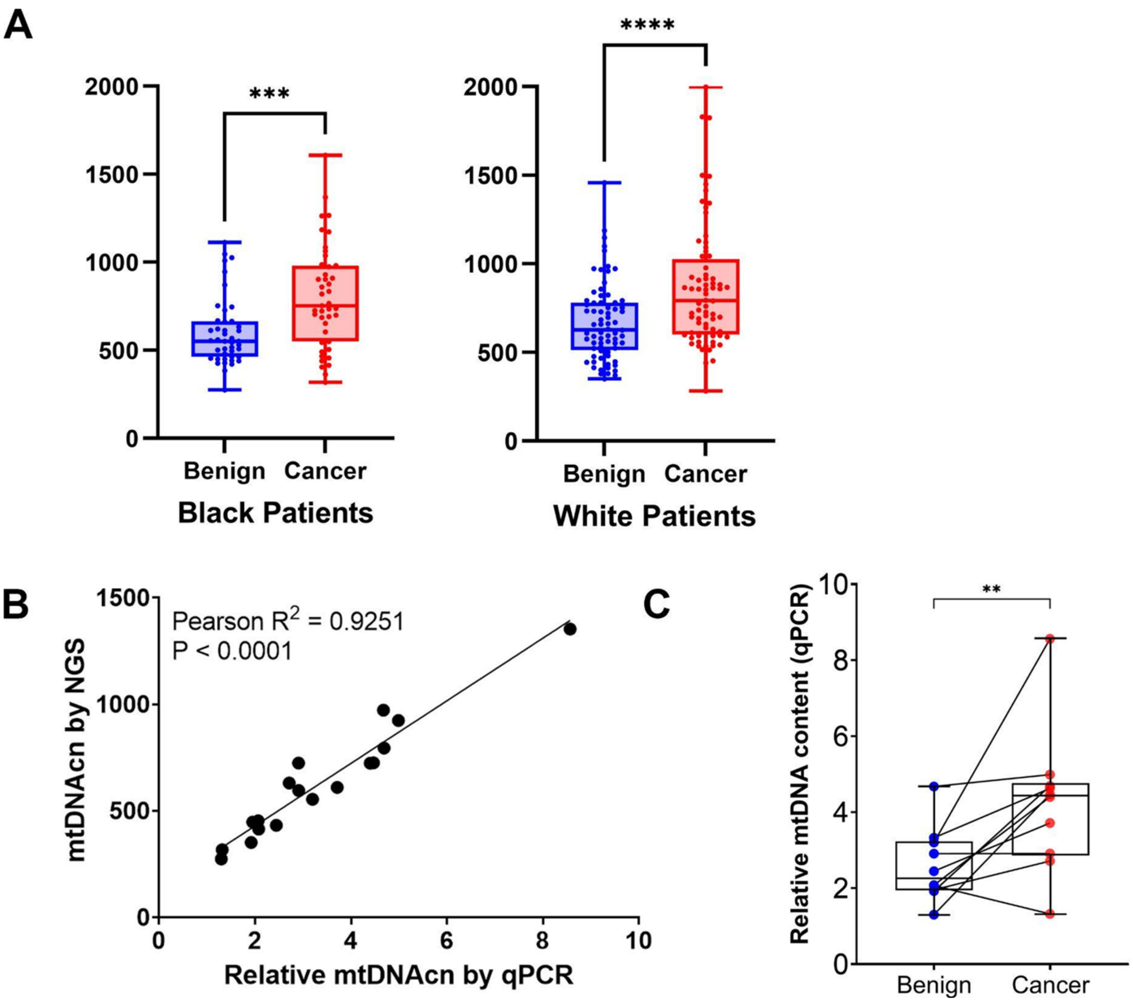


**Supplemental Figure 4**. **Increased mtDNAcn occurs in prostatic adenocarcinoma tissues from both Black and White men.** (**A**) Similar increases in mtDNAcn were found in tumors compared to matched benign prostate samples from Black and White men after WGS. (**B**) Scatter plot shows a strong correlation of mtDNA levels when comparing the results from WGS to qPCR using the same prostate samples from LCM. (**C**) Relative mtDNAcn by qPCR shows a similar pattern of increased levels in tumor tissue to the results found by *in situ* hybridization (Fig. 2) and WGS (Fig. 3) (n = 10 patients). **P < 0.0021, ***P < 0.0002, and ****P < 0.0001.

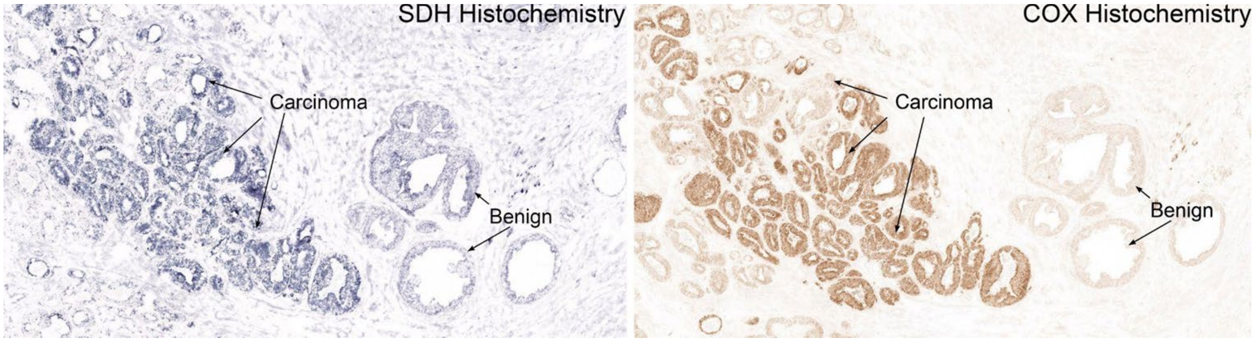


**Supplemental Figure 5. Increased COX and SDH enzyme activity is observed in prostate cancer compared to normal adjacent glands**. (**A**) SDH histochemistry and (**B**) COX histochemistry both reveal increased and heterogenous signals in the cancer regions, similar to the results of mtDNA *in situ* hybridization. Original magnification, x70.

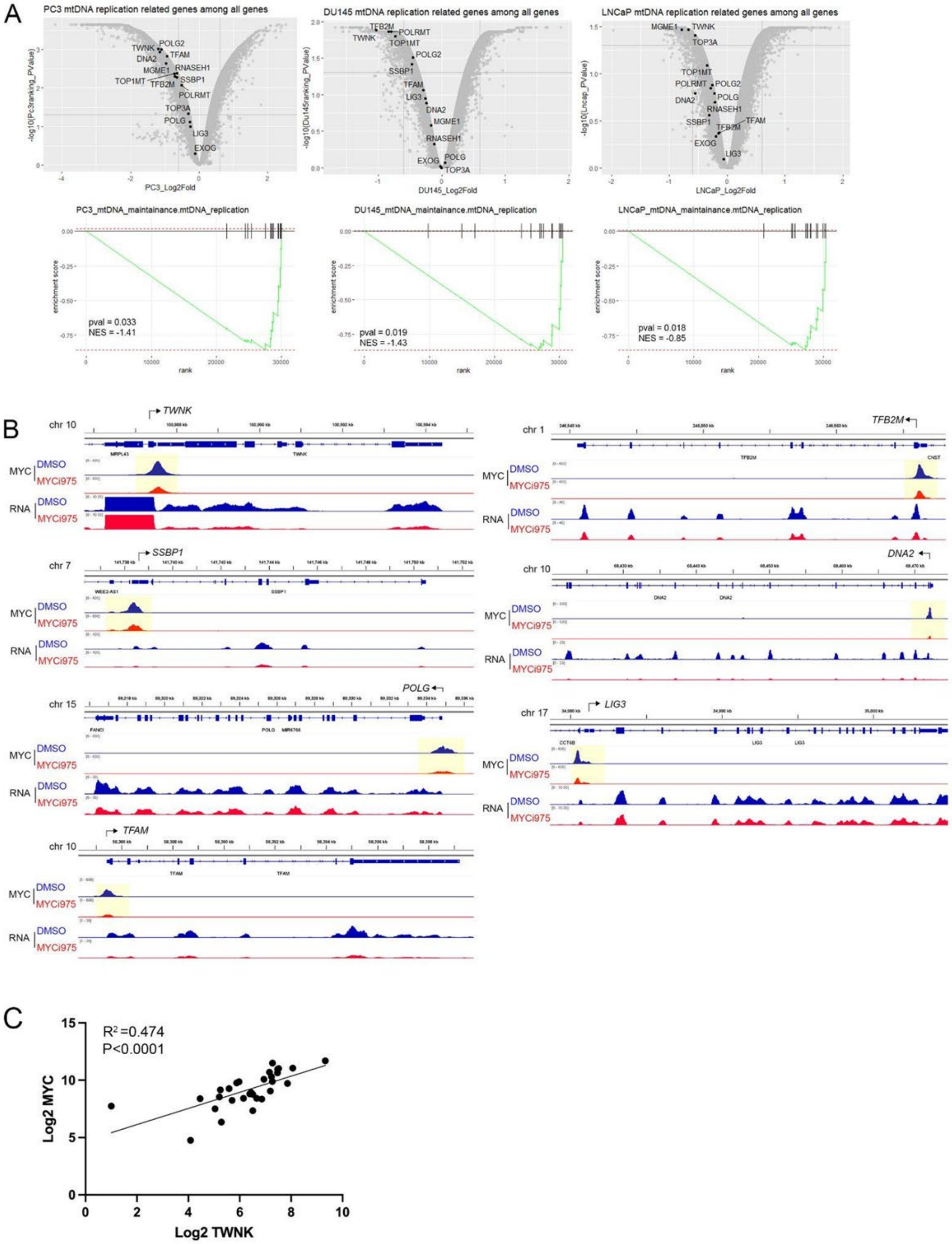


**Supplemental Figure 6. MYC regulates a number of genes encoding factors involved in mtDNA replication.** (**A**) Microarray analysis in cell lines after siRNA-based MYC knockdown (59) reveals down regulation of mtDNA replication-related genes from 3 different human prostate cancer cell lines. (**B**) Publicly available ChIP-seq and RNA-seq data using 22Rv1 prostate cancer cells shows MYC protein binds to the promoter region of genes encoding proteins for mtDNA replication, and MYC occupancy and expression levels were decreased after the cells were treated with a MYC inhibitor. (**C**) Correlation between *MYC* and *TWNK* (encoding TWINKLE) in mCRPC cases in the COMBAT-CRPC study. Each point represents the Log2 mRNA TPM from RNAseq from laser captured frozen tissue biopsy samples.

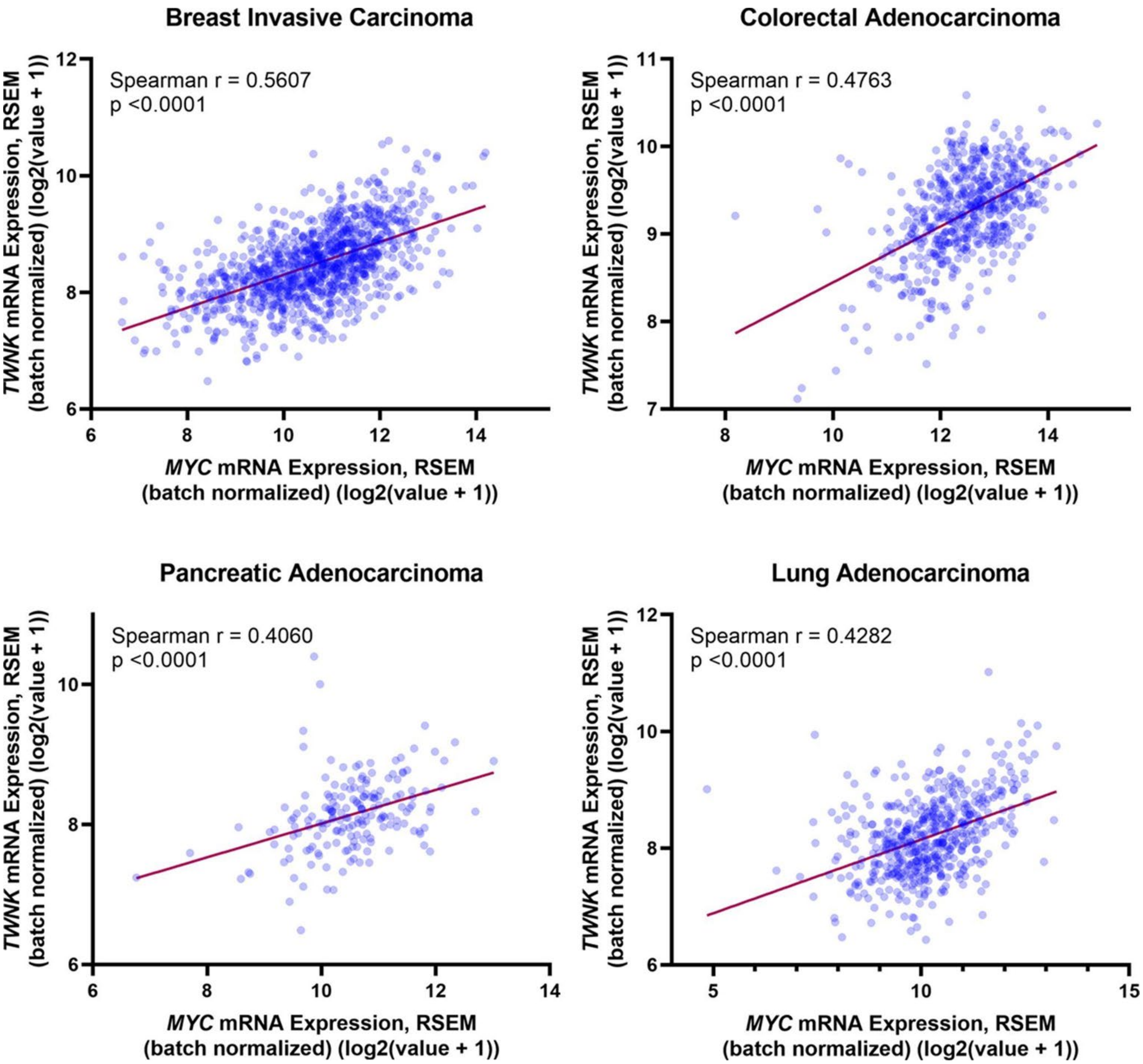


**Supplemental Figure 7. Correlation between *TWNK* and *MYC* RNA expression in various cancer types.** *TWNK* expression level is correlated with *MYC* expression in multiple cancers. Lines indicate the best fit linear trend between these two transcripts.

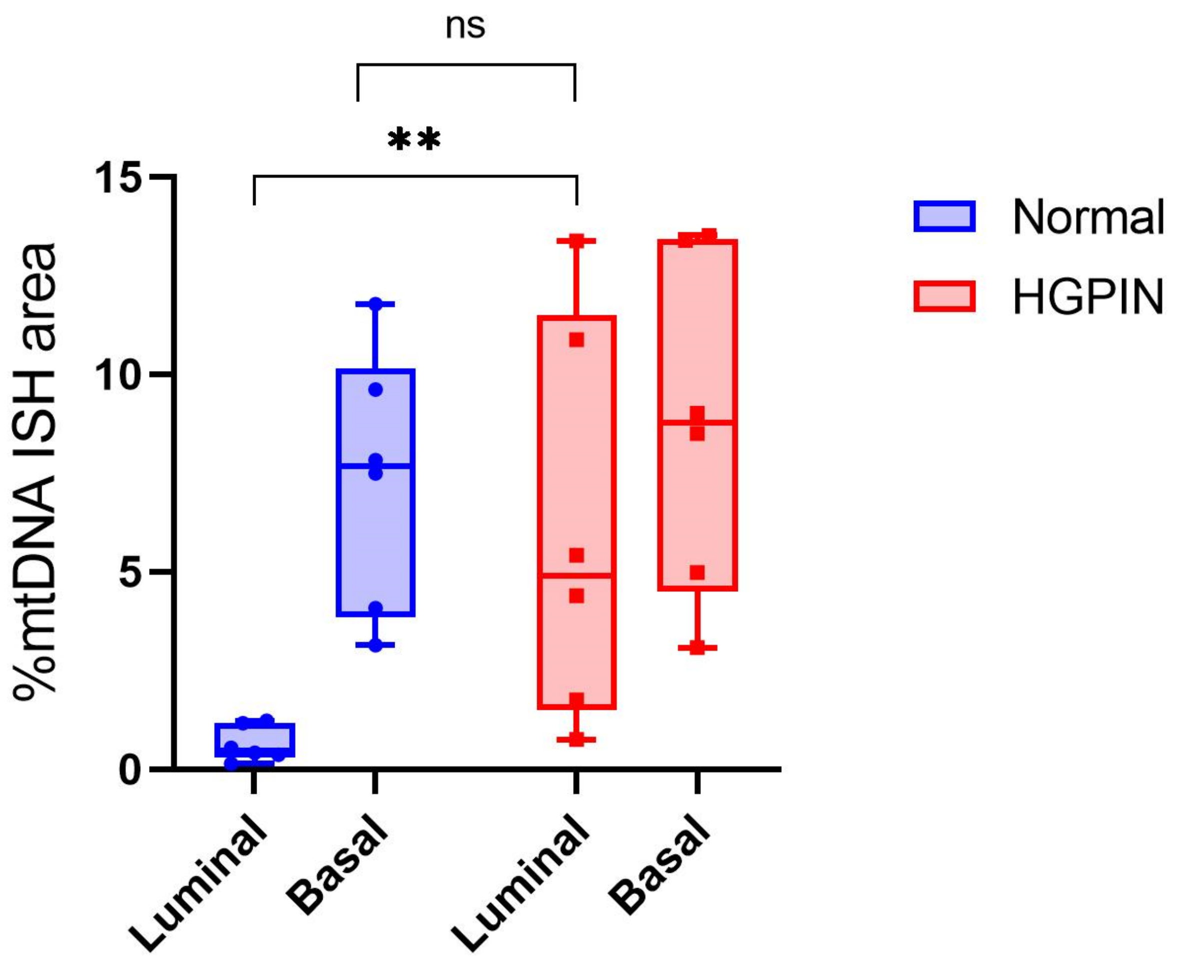


**Supplemental Figure 8. HGPIN luminal cells exhibit higher mtDNA *in situ* hybridization signals than normal luminal cells in the human prostate.** Tissues were subjected to multiplex CISH-IHC as in Fig. 1. Quantitative image analysis results from individual regions of normal and HGPIN glands in which luminal and basal cells were assessed separately. Non-parametric Mann-Whitney tests were done on luminal cells between HGPIN and normal as well as basal cells in normal vs. luminal cells in HGPIN. n = 2 regions per tissue type from 3 patients. **P < 0.0021.

**Supplemental Table 1. Pathology of Cases used in Chromogenic *in situ* hybridization for mtDNAcn.**

| Pt Num | Age Range | Race* | Margins | Grade Group | Gleason Primary | Gleason Secondary | Gleason Sum | Gleason Tertiary | P Stage** |
|---|---|---|---|---|---|---|---|---|---|
| 1 | 60-69 | W | Negative | 2 | 3 | 4 | 7 | | T2N0MX |
| 2 | 60-69 | W | Negative | 2 | 3 | 4 | 7 | | T3AN0MX |
| 3 | 70-79 | B | Positive | 5 | 4 | 5 | 9 | | T3BN0MX |
| 4 | 60-69 | W | Positive | 2 | 3 | 4 | 7 | | T2XNXMX |
| 5 | 60-69 | B | Negative | 3 | 4 | 3 | 7 | 5 | T3AN0MX |
| 6 | 50-59 | W | Positive | 4 | 4 | 4 | 8 | | T3BN0MX |
| 7 | 50-59 | W | Positive | 3 | 4 | 3 | 7 | 5 | T3BN0MX |
| 8 | <50 | W | Negative | 5 | 4 | 5 | 9 | | T3BN1MX |
| 9 | 50-59 | W | Negative | 5 | 4 | 5 | 9 | | T3BN1MX |
| 10 | 70-79 | W | Positive | 3 | 3 | 4 | 7 | | T2N0MX |
| 11 | 50-59 | W | Negative | 3 | 3 | 4 | 7 | | T2N0MX |
| 12 | 60-69 | W | Negative | 3 | 3 | 4 | 7 | | T3AN0MX |
| 13 | 60-69 | W | Negative | 2 | 3 | 4 | 7 | 4 | T2NXMX |
| 14 | 60-69 | W | Positive | 2 | 3 | 4 | 7 | | T2XN0MX |
| 15 | 50-59 | W | Negative | 5 | 4 | 5 | 9 | | T3AN0MX |
| 16 | 60-69 | B | Positive | 2 | 3 | 4 | 7 | | T3ANXMX |
| 17 | 60-69 | W | Positive | 3 | 4 | 3 | 7 | | T3AN0MX |

* Race is self identified. B = Black patients, W = White patients.
** P Stage is the pathological stage at radical prostatectomy using the American Joint Committee on Cancer Staging 2007.

**Supplemental Table 2. Demographic and pathological features of 115 patients with the laser captured microdissected prostate samples for whole genome sequencing and RNAseq.**

| Age | *B | W | Total |
|---|---|---|---|
| **<50** | 3 | 1 | 4 |
| **50-59** | 13 | 28 | 41 |
| **60-69** | 23 | 31 | 54 |
| **70-79** | 4 | 12 | 16 |
| **Total** | 43 | 72 | 115 |

| | ****Pathological Stage** | | | | |
|---|---|---|---|---|---|
| **Grade group** | **T2** | **T3A** | **T3BN0** | **Any T, N1** | **Total** |
| **1** | 9 | 1 | 0 | 0 | 10 |
| **2** | 20 | 14 | 0 | 2 | 36 |
| **3** | 9 | 11 | 7 | 2 | 29 |
| **4** | 3 | 3 | 1 | 0 | 7 |
| **5** | 5 | 11 | 10 | 7 | 33 |
| **Total** | 46 | 40 | 18 | 11 | 115 |

* Race is self identified. B = Black patients, W = White patients.
** P Stage is the pathological stage at radical prostatectomy using the American Joint Committee on Cancer Staging 2007.
